# Connectional axis of individual functional variability: Patterns, structural correlates, and relevance for development and cognition

**DOI:** 10.1101/2023.03.08.531800

**Authors:** Hang Yang, Guowei Wu, Yaoxin Li, Xiaoyu Xu, Jing Cong, Haoshu Xu, Yiyao Ma, Yang Li, Runsen Chen, Adam Pines, Ting Xu, Valerie J. Sydnor, Theodore D. Satterthwaite, Zaixu Cui

**Author notes:** These authors contributed equally (Hang Yang, Guowei Wu). Zaixu Cui **Email:**. **Author Contributions:** Z.C., H.Y., and G.W. designed the study. H.Y. and G.W. performed the analyses with support from Yaoxin Li, X.X., J.C., H.X., and Y.M. J.C. and H.X. replicated all analyses. Z.C. and Yang Li led the acquisition of the YEN dataset, with involvement from X.X., J.C., and H.X. H.Y. created the figures. Z.C. supervised the project. R.C., A.P., T.X., V.S., and T.S. provided feedback on the analyses. H.Y. and Z.C. wrote the manuscript, with review and editing by all other authors. **Competing Interest Statement:** The authors declare no competing interests. **Classification:** Biological Sciences/Neuroscience.

## Abstract

The human cerebral cortex exhibits intricate interareal functional synchronization at the macroscale, with substantial individual variability in these functional connections. However, the spatial organization of functional connectivity (FC) variability across the human connectome edges and its significance in cognitive development remain unclear. Here, we identified a connectional axis in the edge-level FC variability. The variability declined continuously along this axis from within-network to between-network connections, and from the edges linking association networks to those linking the sensorimotor and association networks. This connectional axis of functional variability is associated with spatial pattern of structural connectivity variability. Moreover, the connectional variability axis evolves in youth with an increasing flatter axis slope. We also observed that the slope of connectional variability axis was positively related to the performance in the higher-order cognition. Together, our results reveal a connectional axis in functional variability that is linked with structural connectome variability, refines during development, and is relevant to cognition.

**Significance Statement:** Understanding the spatial organization of inter-individual variability in functional connectivity across the human connectome can offer insights into which connections are most susceptible to insults and interventions. We identify a stable axis on functional connectome along which connectivity variability continuously varies, aligning with structural connectivity variability. We find that the pattern of this axis evolves during development, marked by a decrease in functional connectivity variability among association connections. Moreover, the variability axis is associated with individual differences in higher-order cognitions, driven by increased variability in higher-order association connections linked to enhanced cognitive performance. These findings highlight the importance of connectivity-specific plasticity in understanding both typical and atypical cognitive development across diverse populations.

## Introduction

The human connectome comprises a complex network of interareal connections that facilitates dynamic communication between brain regions. This interregional communication can be represented as functional connectivity (FC) (1, 2), which exhibits substantial organizational differences between individuals(3). This inter-individual variability of functional connectivity is associated with individual differences in human development, cognitive functions, and a wide range of clinical psychopathology (2–5). Recent research has also shown that functional connectivity explains the propagation of neurostimulation-induced signals across the brain and individual variability in functional connectivity helps define the personalized neurostimulation target (6, 7). Accordingly, understanding the spatial patterning of individual variability of functional connections can offer insights into which connections are most susceptible to plasticity, influenced by various factors such as insults and interventions, thereby aiding in the development of connectivity-guided pharmaceuticals and neurostimulation techniques.

Prior studies have examined the spatial organization of inter-individual variability in functional connectivity at the regional level, defined as the summary of connection strengths from one region to all others (3). The regional FC variability is heterogeneously distributed across the cortex and shows a spatially continuous increase along the macroscale cortical axis, which progresses from primary sensorimotor cortices to higher-order association cortices (3, 8, 9).

However, regional-level FC variability, being an aggregate measure, lacks the nuanced and rich information provided by studying variability at the level of individual functional connectivity edge (10, 11). Prior studies on functional connectivity related to development, cognition, psychopathology, and neurostimulation are mainly focused on edge-level connectivity rather than regional connectivity (5, 7, 12–14). Thus, examining the spatial patterning of inter-individual variability in edge-level functional connectivity could offer deeper cognitive and clinical insights than those provided by regional connectivity variability alone. Despite this potential, few studies have systematically analyzed the spatial organization of individual variability in functional connectivity at the edge level across the human connectome.

Previous studies have introduced the concept of an axis to quantify the orderly spatial progression of neurobiological properties that change systematically across the brain (8, 15). In the present study, we aimed to empirically characterize the connectome-wide spatial heterochronous pattern of individual FC variability and hypothesized the presence of a stable, reproducible axis across connectome edges, referred to as the “connectional axis”, along which individual FC variability continuously varies. Additionally, convergent evidence from multiple independent studies has demonstrated that the structural connectivity of the white matter tracts both facilitates and constrains the transfer of functional signals, thereby forming the backbone for the emergence of functional connectivity (1, 16). Previous research has shown a similarity between the spatial patterns of regional FC variability and regional structural connectivity variability across the cortex (17, 18). However, the extent to which this relationship holds at the connectome edge level remains unclear. We hypothesized that the connectional axis of individual FC variability would be associated with structural connectivity variability across connections.

Functional connectivity undergoes significant refinement throughout childhood and adolescence (19–22), with this neurodevelopmental process playing a critical role in shaping the spatial pattern of individual variability in functional connections. Connections among higher-order association regions exhibit prolonged maturation during youth, governed by conserved genetic programs. These include synaptic and axonal pruning of redundant or variable local connections (23), and activity-dependent myelination of evolutionarily conserved long-range pathways (24). While experience-driven plasticity may amplify individual differences, the shared genetically encoded process and environmental demands—such as social interaction and skill learning—could drive convergent optimization of association circuits across individuals. We therefore hypothesize that the development during this period ultimately reduce individual variability in association connectivity as these networks stabilize into efficient, conserved configurations. Here, we provide the first systematic examination of the development of the connectome-wide spatial pattern of FC variability during youth. Additionally, functional connectivity among association networks has been linked to individual differences in higher-order cognition, such as executive function (2, 25–27). However, it remains unclear how the connectome-wide spatial organization of individual FC variability relates to cognitive performance. Prior work showed that greater neural flexibility is associated with better higher-order cognitive abilities (27–29). Therefore, we hypothesized that populations with greater individual variability in association connectivity would exhibit superior higher-order cognitive abilities (e.g., executive function) compared to those with lower variability.

To address these hypotheses, we utilized functional MRI (fMRI) data from three independent, high-quality datasets to non-invasively study the spatial organization of individual FC variability across the human connectome. FC was defined as the synchronization of the blood oxygen level-dependent (BOLD) fMRI signals between two brain regions. Prior research has defined the brain axes or gradients by ranking the magnitudes of neurobiological properties (30, 31). Following a similar methodology, we defined a connectional axis of individual FC variability across all edges of the cortical connectome. Our findings reveal that the connectional axis of FC variability extends from within-network connections to between-network connections, and from connections linking association networks to those linking sensorimotor networks. Interestingly, connections between association and sensorimotor networks exhibited the lowest FC variability, rather than the connections within sensorimotor networks. Furthermore, we discovered that the pattern of FC variability along the connectional axis is associated with the spatial pattern of structural connectivity variability. Lastly, our results indicate that the connectional variability axis is refined during youth, and associates with higher-order cognitive functions. All primary findings were established using developmental and young adult datasets from the Human Connectome Project (32, 33), and subsequently replicated in an independent cohort of Chinese youth.

## Results

We harnessed two independent datasets, namely the Human Connectome Project (HCP)-development (HCP-D, *n* = 486, 265 females, aged 8–21 years) (33) and unrelated HCP-young adult (HCP-YA, *n* = 275, 146 females, aged 22–35 years) (32), to estimate the inter-individual variability of functional connectivity (FC). To this end, we included 26- and 58-min blood oxygen level-dependent (BOLD) functional MRI data from the HCP-D and HCP-YA datasets, respectively. Furthermore, we included a Chinese cohort, the Youth Executive function and Neurodevelopment (YEN, *n* = 218, 113 females, aged 6–22 years), as a replication dataset. This cohort consists of 18-min BOLD functional MRI data for each participant. FC was calculated for each participant based on the a priori Schaefer cortical parcellation atlas of 400 regions (34). We determined FC as the Pearson correlation coefficients between pairs of regional time series, resulting in a 400×400 symmetrical FC matrix for each participant. The Pearson correlation coefficients were then *r*-to-*z* transformed using the Fisher transformation. This full FC matrix is termed a functional connectome. The matrix contained 79,800 unique elements, and we denoted each element as an “edge” connecting two cortical regions. A linear mixed-effects model was applied to estimate the inter- and intra-individual variability of each FC edge (35). As in previous studies (3, 36), we used the intra-class coefficient (ICC) to measure the adjusted inter-individual variability while accounting for intra-individual variability. Finally, a 400×400 symmetrical inter-individual variability matrix was generated, with each element representing the variation in the connection strength for an edge across all participants. We grouped the 400 cortical regions into networks using the Yeo atlas (37), which consists of visual, somatomotor, dorsal attention, ventral attention, frontoparietal, default mode, and limbic networks. In general, the visual and somatomotor networks are assigned to sensorimotor networks, and the other five networks belong to association networks. However, the limbic network was excluded from subsequent analyses because of a substantial signal loss, especially in the orbitofrontal and ventral temporal cortex(38). Finally, we evaluated 374 cortical regions with 69,751 unique edges.

### Individual variability in edge-level FC progressively varies along a stable and reproducible connectional axis

We arranged the 374 cortical regions based on their affiliations to the large-scale functional networks from priori Yeo atlas(37) (**Fig. 1a**), and generated matrices of individual FC variability for both the HCP-D (**Fig. 1b**) and HCP-YA (**Fig. 1c**) datasets. Visual inspection suggested that individual variability was heterogeneously distributed across connectome edges in both datasets. Specifically, high variability was prevalent in the connections within or between association networks, including the default mode, frontoparietal, ventral attention, and dorsal attention networks. In contrast, we observed lower individual variability in the between-network edges bridging the association and sensorimotor regions. We found a significant correlation (Spearman’s rho = 0.916, *P*_perm_ < 0.0001, confidence interval [CI] = [0.915, 0.918], two-sided) between the individual variability of the HCP-YA and HCP-D datasets across all edges, indicating a highly stable pattern of edge-wise individual variability across datasets.

**Figure 1.**
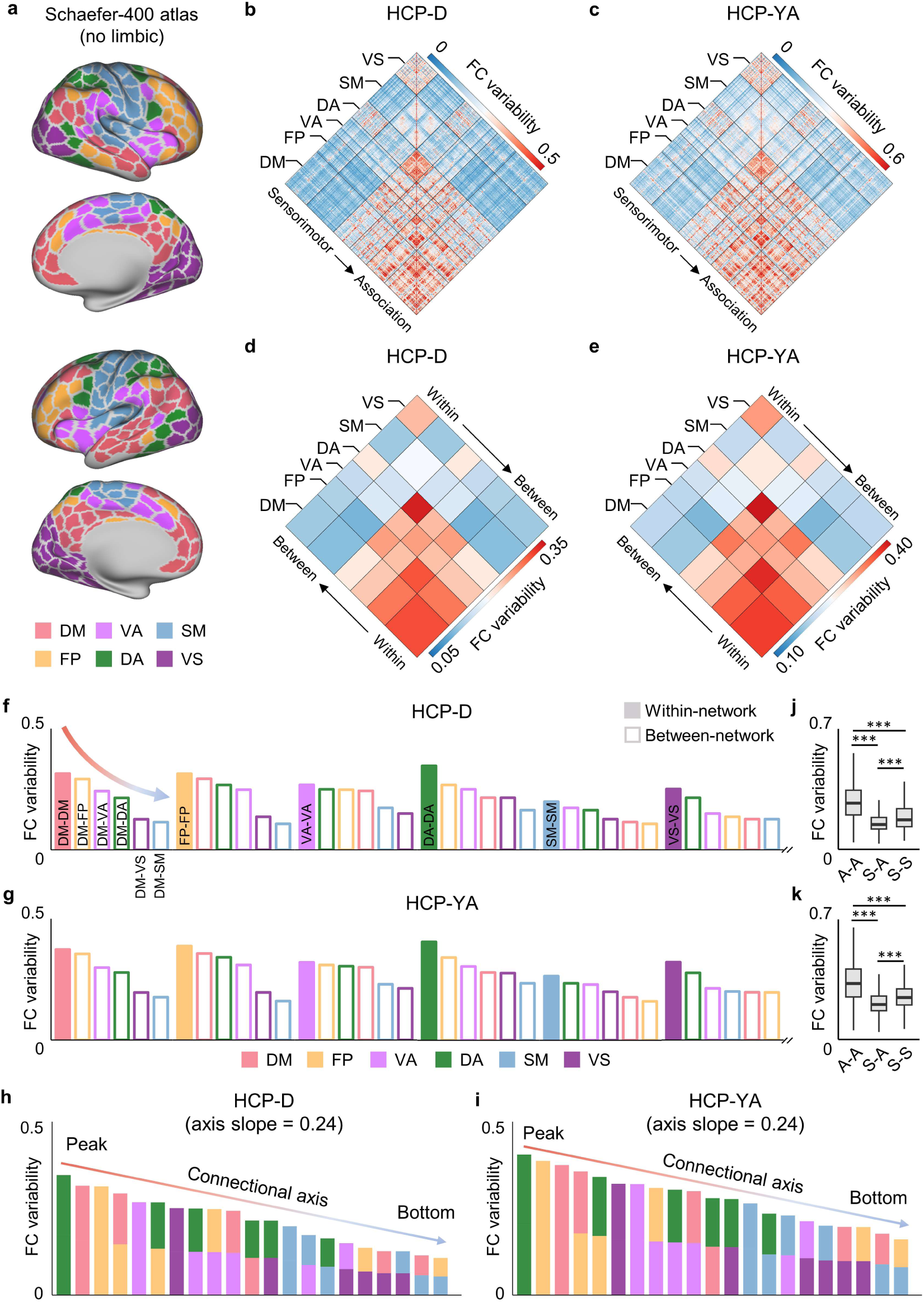
Individual variability in edge-level FC declines along a connectional axis. **a**, The Schafer-400 atlas excludes the limbic network. The cortical regions are assigned to each of the six large-scale functional networks from priori Yeo atlas, which include the default mode (DM), frontoparietal (FP), ventral attention (VA), dorsal attention (DA), somatomotor (SM), and visual (VS) networks. **b**, Inter-individual FC variability across all participants at the edge level in the HCP-D dataset. Individual variability was heterogeneously distributed across the functional connectome edges. **c**, Inter-individual FC variability across all participants at the edge level in the HCP-YA dataset. **d**,**e**, Network-level average individual FC variability indicates that for each association network, the FC variability continuously declines from within-network to between-network connections for the HCP-D (**d**) and HCP-YA (**e**) datasets. **f**,**g**, Bar plots represent the connectional variability axis for each network separately in the HCP-D (**f**) and HCP-YA (**g**) datasets. For each network, individual FC variability declines from within-network to between-network edges along a continuous axis. **h**,**i**, Ordering the individual variability across all the 21 network-level edges reveals a continuous axis from edges within and between association networks to the edges between association and sensorimotor networks for HCP-D (**h**) and HCP-YA (**i**) datasets. The variabilities of connections within networks and that between association networks were above the dotted line, whereas the variabilities of connections between association and sensorimotor networks were below the dotted line. **j**,**k**, Edge-wise individual variability within association networks (A-A) was significantly higher than those within sensorimotor networks (S-S) and between sensorimotor and association networks (S-A) in HCP-D (**j**) and HCP-YA (**k**) datasets. Additionally, S-S connections exhibit higher variability than S-A connections in both datasets. A permutation test with 10,000 iterations was applied to test the group differences in average FC variability. *** indicates *P*_perm_ < 0.0001. FC, functional connectivity; HCP, Human Connectome Project; HCP-D, HCP development; HCP-YA, HCP young adult.

Next, we grouped the cortical regions into six major networks and averaged FC variabilities within and between the networks, generating network-level 6×6 matrices for the HCP-D (**Fig. 1d**) and HCP-YA (**Fig. 1e**) datasets. Prior studies have consistently defined a brain axis or gradient by ranking the neurobiological properties from the highest to the lowest (30, 31). With this approach, we observed an axis of decline in inter-individual FC variability progressing from within-network connections to between-network connections for each large-scale network. For example, individual variability of the default mode network declined along an axis progressing from edges within the default mode network to those between the default mode and other association networks, further decreasing at the edges between the default mode and sensorimotor networks. We termed these axes of individual variability as the “connectional variability axis” for each network. These network-level connectional axis patterns were highly consistent between the two independent datasets (Spearman’s rho = 0.987, *P* = 4.45×10^-6^, CI = [0.937, 0.999], two-sided).

We separately ranked within-network and between-network variabilities for each network in the HCP-D (**Fig. 1f**) and HCP-YA (**Fig. 1g**) datasets to intuitively visualize the connectional variability axis. We found that the ranking for each network was highly consistent between HCP-D and HCP-YA, suggesting the robustness and reproducibility of the connectional variability axis. The ranking demonstrated that for each association network, individual variability declined from within-network edges to edges between two association networks and further declined for connections between the sensorimotor and association networks.

Having delineated the connectional variability axis for each association network, we examined the whole-brain connectome. We ranked the inter-individual variability of all 21 within-network and between-network connections for the HCP-D (**Fig. 1h**) and HCP-YA (**Fig. 1i**) datasets. The connections within the association networks were at the peak of this axis, whereas the between-network edges connecting the sensorimotor and other networks were at the base. In particular, we introduced an axis slope to measure the heterogeneity of connectional variability across the connectome. The axis slope was quantified by performing a linear regression on the 21 ranked network-level variability values. The estimated axis slope is 0.24 for both HCP-D and HCP-YA.

Then, we categorized all the 69,751 unique connections into three groups: within-association network edges (A-A), within-sensorimotor network edges (S-S), and between sensorimotor and association network edges (S-A). A permutation test of 10,000 iterations, permuting the edges across the groups, was performed to test for differences in variability levels between the three groups. We found that the individual variability of A-A connections was significantly higher than those of S-A and S-S connections (*P*_perm_ < 0.0001) in the HCP-D (**Fig. 1j**, Cohen’s d = 0.95, A-A vs. S-S; Cohen’s d = 1.44, A-A vs. S-A) and HCP-YA (**Fig. 1k**, Cohen’s d = 0.77, A-A vs. S-S; Cohen’s d = 1.26, A-A vs. S-A) datasets. Furthermore, S-S connections displayed significantly higher variability than S-A connections (*P*_perm_ < 0.0001) in both the HCP-D (Cohen’s d = 0.43) and HCP-YA (Cohen’s d = 0.49) datasets.

### The connectional axis pattern of FC variability is associated with the individual variability of structural connectivity communicability across connectome edges

Previous studies have reported that structural connectivity constrains the dynamic communication between cortical regions and shapes FC patterns (1, 39), suggesting that structural connectivity variability can shape FC variability. Notably, direct structural connections alone do not capture the full spectrum of functional interactions, as FC can emerge from indirect or polysynaptic communication. However, the polysynaptic interactions between structurally unconnected brain areas can be measured using communicability (40, 41). Communicability is derived from the edge-level structural connectivity and quantifies the potential of two regions to communicate through other regions via pathways of all possible topological lengths (41). A recent study has demonstrated that communicability provides the most accurate and robust accounts of stimulus propagation through white matter connectivity (40). We hypothesized that inter-individual variability in structural connectome-based communicability would be associated with inter-individual variability in FC across connectome edges.

We used diffusion MRI datasets to reconstruct whole-brain white matter tracts of individual participants via probabilistic fiber tractography with multi-shell, multi-tissue constrained spherical deconvolution (42). Anatomically constrained tractography (ACT) (43) and spherical deconvolution informed filtering of tractograms (SIFT) (44) were applied to improve the biological accuracy of fiber reconstruction. The reconstructed fiber tracts were visually inspected; see **Fig. S1** for exemplar visualizations for four randomly selected participants.

We quantified the number of streamlines connecting every pair of cortical regions as defined by the Schaefer atlas, resulting in a structural connectome of streamline counts for each participant (**Fig. 2a**). We then calculated the communicability matrix of each participant’s structural connectome for the HCP-D and HCP-YA datasets, respectively, which provided a communicability measure for each edge. We acquired an inter-individual variability matrix of structural communicability for each dataset by evaluating the mean absolute deviation across all participants and log-transformed the variability matrices.

**Figure 2.**
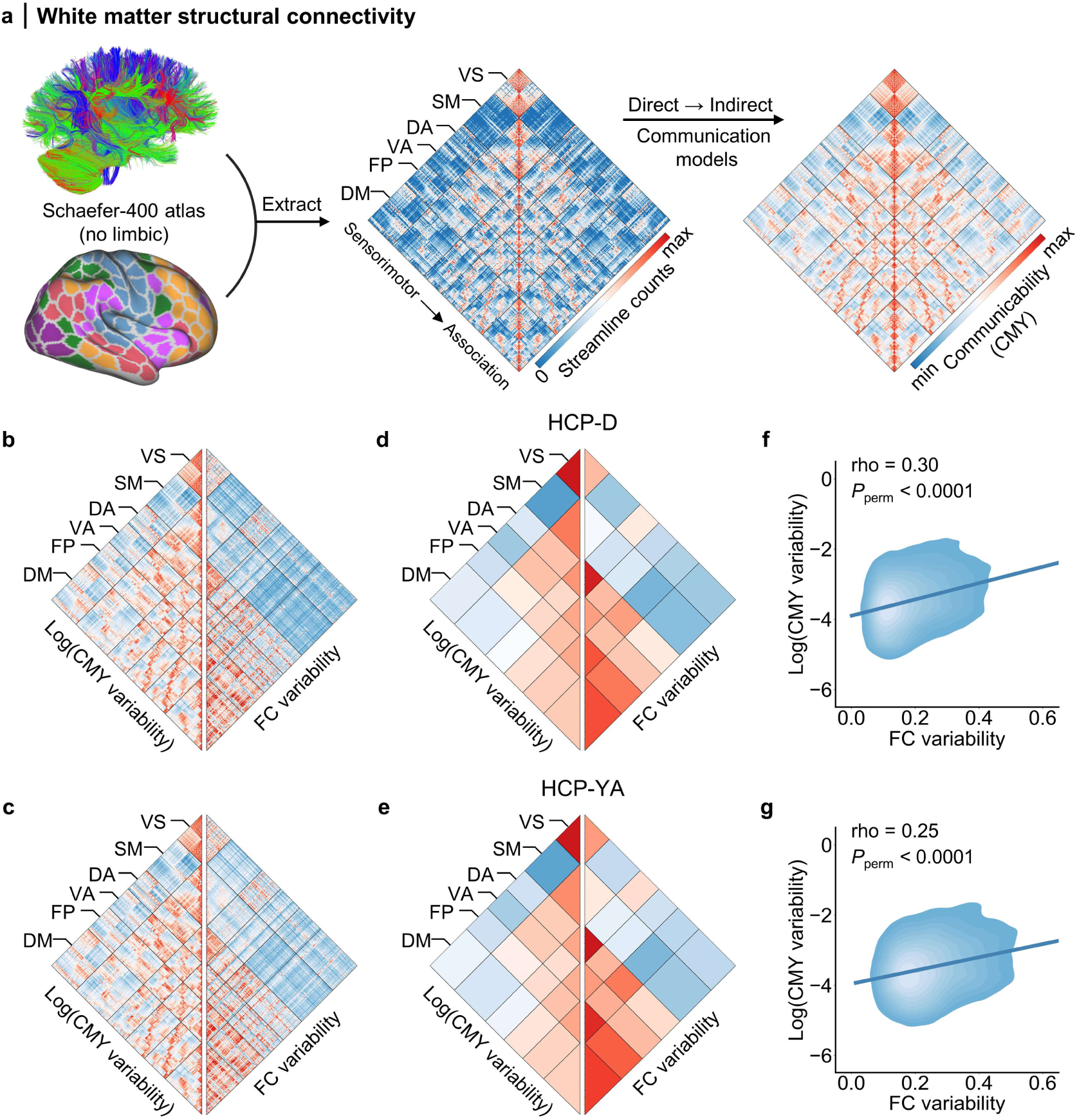
Individual variability in structural connectivity communicability is associated with the connectional axis pattern in FC variability across connectome edges. **a**, Structural connectomes were constructed using white matter tracts tractography from diffusion MRI data and the Schaefer atlas. Edge strengths in the structural connectome indicate the white matter fiber density between pairs of regions. The communicability matrix was derived from structural connectivity to account for the indirect communication between regions without direct structural connections in both datasets. **b**,**c**, Side-by-side comparison of the matrices for individual variability of structural communicability and individual FC variability in the HCP-D (**b**) and HCP-YA (**c**) datasets. The communicability variability was log-transformed. **d**,**e**, Network-level average of structural communicability variability and FC variability in HCP-D (**d**) and HCP-YA (**e**). **f**,**g**, Individual FC variability was positively correlated with individual variability of structural communicability across all edges in both HCP-D (**f**, Spearman’s rho = 0.30, *P*_perm_ < 0.0001, CI = [0.29, 0.31], two-sided) and HCP-YA (**g**, Spearman’s rho = 0.25, *P*_perm_ < 0.0001, CI = [0.24, 0.25], two-sided) datasets. A permutation test was performed 10,000 iterations by randomly shuffling the rows and columns of the FC variability matrix, then a null distribution was built by evaluating the Spearman’s rank correlation between random FC variability matrices and the true structural communicability matrix. CMY, communicability.

Similar to the edge-level FC variability, we observed that individual variability in structural communicability was heterogeneously distributed across connectome edges. To facilitate comparison, we showed the upper triangle of the structural communicability variability and individual FC variability matrices side by side for the HCP-D (**Fig. 2b**) and HCP-YA (**Fig. 2c**) datasets. Network-level averages demonstrated a similar pattern between structural communicability variability and FC variability for both datasets (**Fig. 2d** and **Fig. 2e**). Individual variability in structural communicability declined from connections among association networks to those between association and sensorimotor networks, similar to that of FC variability. For example, structural communicability variability declined from the edges between the default mode network and other association networks to those between the default mode and sensorimotor networks. Finally, we found a significant correlation between FC variability and structural communicability variability across all edges for the HCP-D (Spearman’s rho = 0.30, *P*_perm_ < 0.0001, CI = [0.29, 0.31], two-sided; **Fig. 2f**) and HCP-YA (Spearman’s rho = 0.25, *P*_perm_ < 0.0001, CI = [0.24, 0.25], two-sided; **Fig. 2g**) datasets. This finding indicates that edges with higher individual variability in structural communicability showed relatively higher variability in FC. These results provide a potential structural basis for the observed pattern of connectional axis in individual FC variability.

### Connectional variability axis evolves in youth with a more uniform individual FC variability across connectome edges

Prior studies have shown that age-related development contributes to variations in FC among individuals, and the neurodevelopmental timing of functional connections varies according to their spatial position in human connectome (8, 24, 45, 46). Therefore, we hypothesized that the connectional axis pattern of FC variability would evolve during youth, a critical period for the development of higher-order cognition.

To test this hypothesis, we examined participants (aged 8–21) from the HCP-D dataset. We used a sliding-window approach to sort the participants by age and divided them into groups (**Fig. 3a**). The window length was set to 50 participants, with a step size of five participants. This procedure resulted in 88 groups with progressively increasing age ranges. For each group, we computed an individual variability matrix for edge-level FC across all the participants (**Fig. 3b**). A group-average age and a group-average head motion were assigned to each group. The sex of each group is represented as the percentage of male participants.

**Figure 3.**
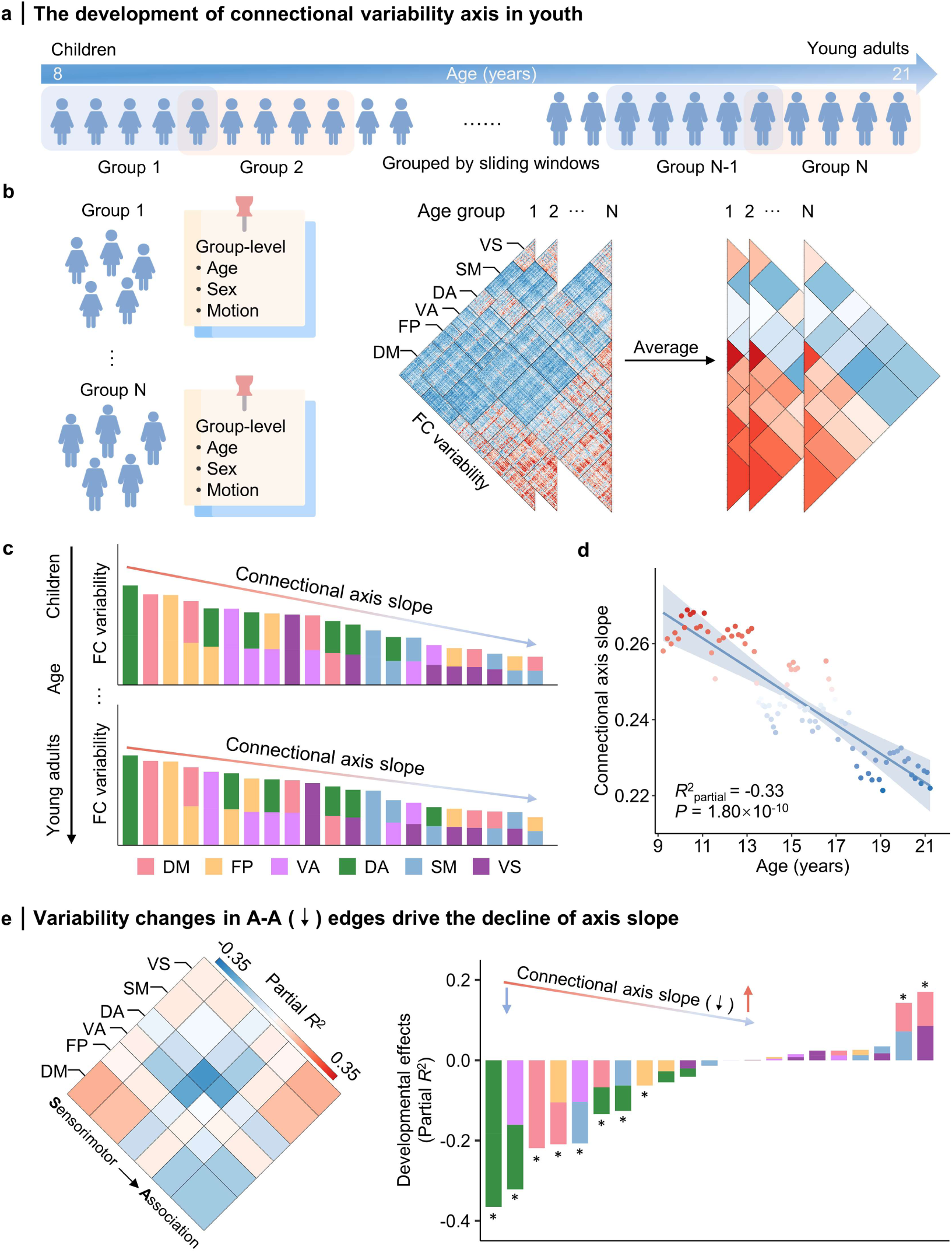
Connectional axis of FC variability evolves during youth. **a**, Participants in the HCP-D dataset were ordered based on age and subsequently divided into 88 groups using a sliding window approach. The window size and step were set to 50 and 5 participants, respectively. **b**, Inter-individual FC variability matrices were computed for each of the 88 groups within the HCP-D dataset. **c**, Network-level average FC variability was arranged in descending order to evaluate the axis of network-level variability and the slope of the variability axis was fitted for each group. **d**, A GAM analysis indicated that the slope of the connectional variability axis declined significantly with age in youth, suggesting that the pattern of FC variability becomes more uniform across connectome edges during youth. **e**, The decline of FC variability in association-to-association (A-A) edges mainly drive the decline of variability axis slope during development. GAM analysis was performed for FC variability of the 21 network-level connections separately. The asterisk (*) indicates *P* < 0.05 (FDR adjusted). VS, visual; SM, somatomotor; DA, dorsal attention; VA, ventral attention; FP, frontoparietal; DM, default mode.

We evaluated how the connectional axis of the individual FC variability changed during youth. For each of the 88 groups, we computed the network-level average of individual FC variability. Then, we ranked the resulting 21 network-level variability values from the highest to the lowest for each group, respectively (**Fig. 3c**), as in **Fig. 1h, i**. Specifically, we observed that the axis slope for the 21 network-level variability differs between children and young adults. An increase in the axis slope could indicate an increased difference between FC variabilities of connections among association networks at the top end of the connectional variability axis and that of sensorimotor-association connections at the bottom end, whereas a decrease in the axis slope could indicate a reduced difference between them. Using GAM analysis, we found that the slope of the network-level FC variability axis significantly declined during youth (partial *R*^2^ = -0.33, *P* = 1.80×10^-10^, **Fig. 3d**). This result suggests that individual FC variability becomes more uniform across connectome edges during development.

We further examined the driving factors underlying the developmental changes of the connectional axis slope of FC variability. To do this, we evaluated the changes in the FC variability for each of the 21 network-level connections. We found that the FC variability of 10 connections was statistically significant (*P*_FDR_ < 0.05, **Fig. 3e**). Particularly, the connections within the dorsal attention network, between dorsal and ventral attention networks, and within the default mode network exhibited the most pronounced decline of FC variability with age. In contrast, connections between the default mode network and visual/somatomotor network showed an increase in FC variability, though the effect size was limited. These results suggest that the prominent reduction in FC variability in association-to-association (A-A) connections primarily drives the developmental changes of the connectional axis slope of FC variability.

### Connectional variability axis is associated with individual differences in higher-order cognition

Having found that the connectional axis of FC variability evolves during youth, we evaluated the implications of this variability axis on cognition. Higher-order cognitive functions, such as working memory and inhibition, exhibit prolonged development during youth and show significant individual differences in adulthood. Here, we examined the relationship between the connectional variability axis and individual differences in higher-order cognitive functions using the HCP-D and HCP-YA datasets. The composite score of fluid cognition provided by the datasets summarizes a broad range of higher-order cognitive functions, including flanker inhibition, dimensional change card sort (flexibility), picture sequence memory, list sorting working memory, and pattern comparison processing speed (47).

The HCP-D dataset included 352 participants, whereas the HCP-YA dataset contained 274 unrelated participants with fluid cognition scores. As in the developmental analysis, we sorted all participants based on their cognitive performance and divided them into groups using a sliding-window approach for both datasets (**Fig. 4a**). Using a window size of 50 participants and a step size of 5 participants, we obtained 61 groups for the HCP-D and 45 groups for the HCP-YA. We computed a matrix of individual FC variability across all participants for each group (**Fig. 4b**). The ranking of network-level average FC variability indicated varying slope values for the variability axis across the different cognitive groups (**Fig. 4c**). After controlling for age, sex, and in-scanner motion, the GAM analysis revealed that a higher fluid cognition score was significantly associated with a steeper slope in the connectional variability axis in both the HCP-D (*t* = 3.53, *P* = 8.41×10^-4^, partial *R*^2^ = 0.15, **Fig. 4d**) and HCP-YA (*t* = 5.08, *P* = 9.62×10^-6^, partial *R*^2^ = 0.41, **Fig. 4e**) datasets. These results suggest that groups of participants with stronger higher-order cognition displayed a steeper slope in the connectional axis, indicating a greater difference between among-association variabilities at the top end of the axis and sensorimotor-association variabilities at the bottom end.

**Figure 4.**
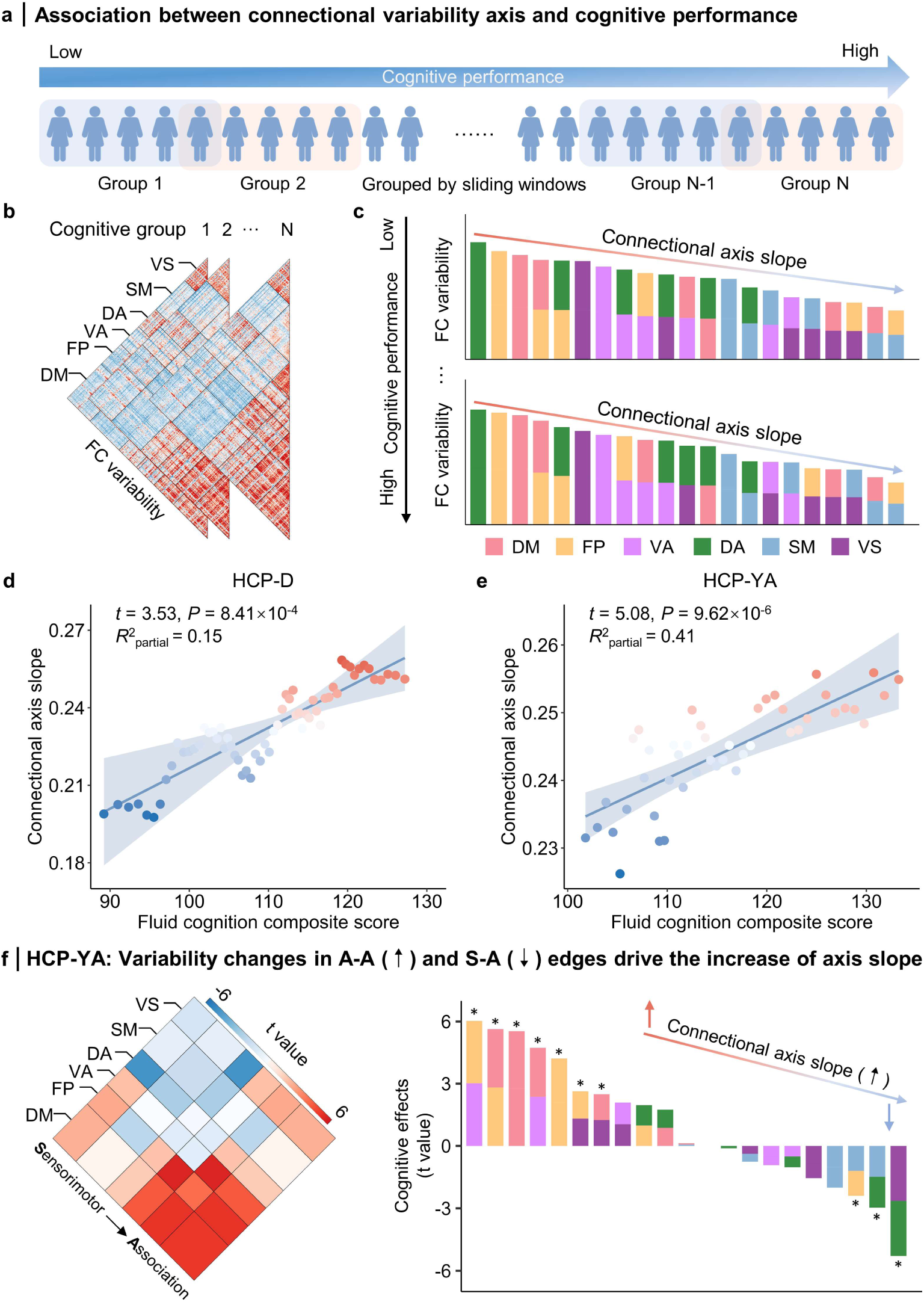
The connectional variability axis pattern is associated with the individual differences in higher-order cognitive functions. **a**, Participants were sorted based on their cognitive performance and divided into groups using a sliding window approach, with a window and step size of 50 and 5 participants, respectively. A total of 61 groups and 45 groups were obtained for the HCP-D and HCP-YA datasets, respectively. **b**, The matrix of individual FC variability was computed across all participants for each cognitive subgroup. **c**. Illustration of changes in the connectional axis slope of FC variability with different higher-order cognitive performance. A linear regression was performed to acquire the slope of the variability axis for each group. **d**,**e**, The connectional axis slope of FC variability showed a significant positive association with the fluid cognition composite score in both the HCP-D (**d**) and HCP-YA (**e**) datasets. Each data point represents a specific group in the analysis. **f**, In the HCP-YA dataset, the increase of FC variability in association-to-association (A-A) edges and decline of sensorimotor-to-association (S-A) variability drives the increase of axis slope with better higher-order cognition. GAM analysis was performed for FC variability of the 21 network-level connections separately. The asterisk (*) indicates *P* < 0.05 (FDR adjusted). VS, visual; SM, somatomotor; DA, dorsal attention; VA, ventral attention; FP, frontoparietal; DM, default mode.

We further examined the driving factors underlying the cognitive effects of the connectional variability axis slope. In the HCP-YA dataset, by analyzing the association between FC variability and cognitive performance for each of the 21 network-level connections, we found that connections between the ventral attention network and the frontoparietal/default mode networks, as well as connections both within and between the default mode and frontoparietal networks, showed a significant (*P*_FDR_ < 0.05) increase in FC variability with better cognitive performance (**Fig. 4f**). In contrast, the FC variability of connections linking dorsal attention and visual/somatomotor networks was negatively related to the cognitive performance. These results suggest that the increased FC variability in association-to-association (A-A) connections and the declined variability in sensorimotor-to-association (S-A) mainly drives the cognition-related changes of the connectional axis slope of FC variability.

### Sensitivity analyses

We conducted sensitivity analyses to evaluate the robustness of our findings to methodological variations and confounding factors. Specifically, we evaluated whether our results remained similar under conditions including, 1) controlling for the influence of FC strength, 2) incorporating the inter-regional distance as a covariate to minimize the effect of spatial proximity, 3) replicating with another widely used HCP multi-modal parcellation (48), and 4) replicating with global signal regression excluded from fMRI preprocessing. Our results indicated that these variations did not change our results regarding the association between FC and SC variability (**Fig. S2**) or developmental and cognitive associations with the connectional variability axis (**Fig. S3**). Additionally, we validated that FC variability also exhibits a significant correlation with variability in structural communicability even when communicability is derived from binarized structural connectomes (**Supplementary Table S1**). Finally, we confirmed that our developmental and cognitive results remain largely consistent regardless of the choice of window length and step size in our sliding window configurations (**Supplementary Table S2-S3**).

### Replication in an independent youth dataset

We replicated all primary findings using an independent Chinese youth sample from the Youth Executive function and Neurodevelopment (YEN) study. Specifically, FC variability was first estimated at the regional level and then averaged at the network level (**Fig. 5a**). The 21 network-level edges were ranked from highest to lowest, forming the connectional axis of the FC variability, with an estimated slope of 0.30 (**Fig. 5b**). A-A connections exhibited the highest individual variability, followed by S-S and S-A connections (**Fig. 5c**). Moreover, individual FC variability was positively correlated with individual variability of structural communicability across all edges (**Fig. 5d**, Spearman’s rho = 0.33, *P*_perm_ < 0.0001, CI = [0.32, 0.34], two-sided). Finally, we found that the slope of the connectional variability axis declined significantly with age in youth (**Fig. 5e**, partial *R^2^* = -0.17, *P* = 0.014) and was positively correlated with executive function scores (**Fig. 5f**, *t* = 3.52, *P* = 0.002, partial *R^2^* = 0.39). See Supplementary Materials for the details of data description.

**Figure 5.**
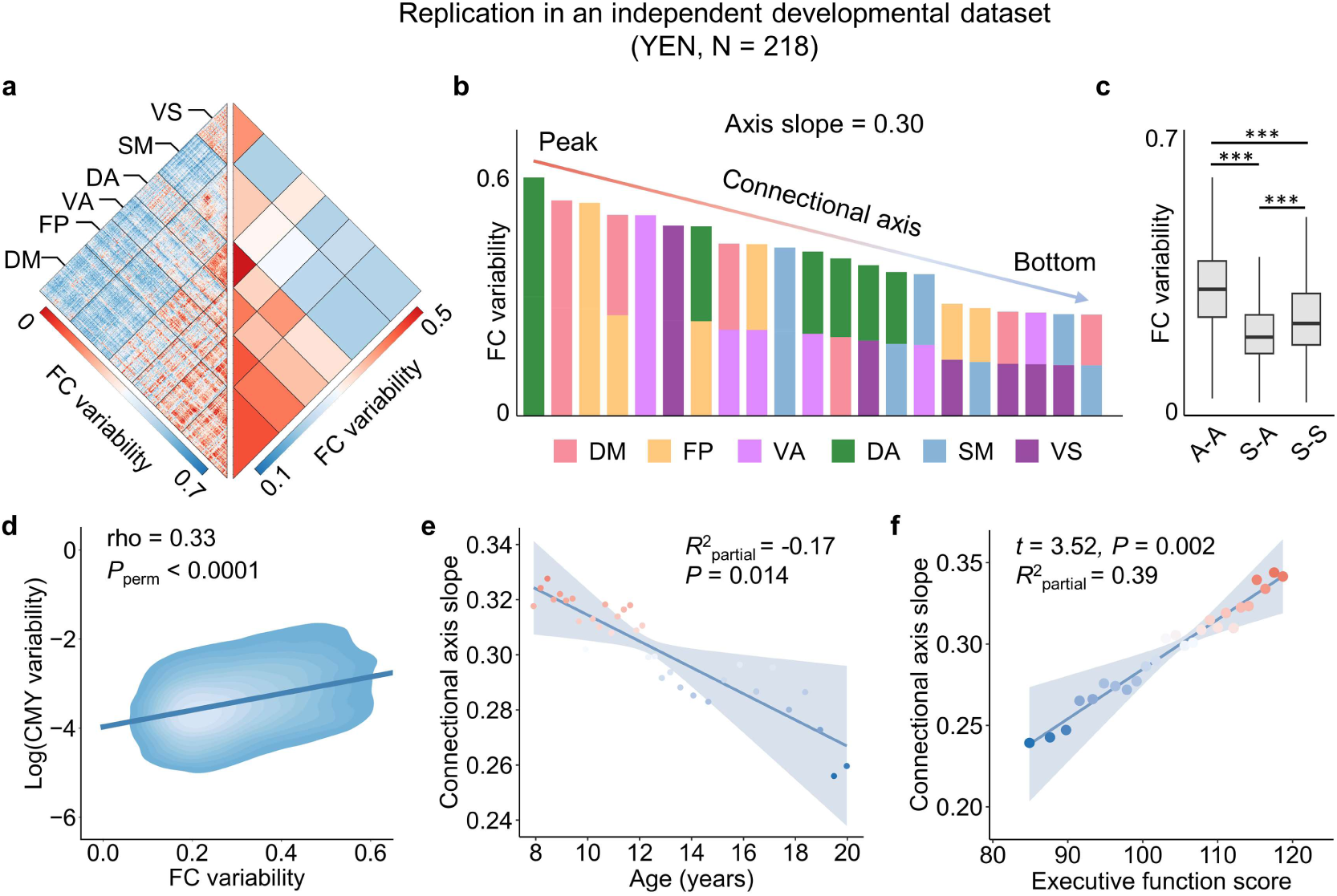
Replication in an independent developmental dataset. **a**, FC variability was estimated across participants from the Youth Executive function and Neurodevelopment (YEN) study at the regional level and subsequently averaged at the network level. The YEN study includes 218 participants, with 113 females, aged 6 to 22 years. **b**, The connectional axis of FC variability is represented by ordering the individual variability across all the 21 network-level edges (axis slope = 0.30). **c**, Edge-wise individual variability was highest within association networks (A-A), followed by those within sensorimotor networks (S-S), and between sensorimotor and association networks (S-A). A permutation test with 10,000 iterations was applied to test the group differences in average FC variability. *** indicates *P*_perm_ < 0.0001. **d**, Individual FC variability was positively correlated with individual variability of structural communicability across all connections (Spearman’s rho = 0.33, *P*_perm_ < 0.0001, CI = [0.32, 0.34], two-sided). **e**, A GAM analysis indicated that the slope of the connectional variability axis declined significantly with age in youth (partial *R^2^* = -0.17, *P* = 0.014). **f**, A significant positive correlation exists between executive function score and the connectional axis slope of FC variability (*t* = 3.52, *P* = 0.002, partial *R^2^* = 0.39). VS, visual; SM, somatomotor; DA, dorsal attention; VA, ventral attention; FP, frontoparietal; DM, default mode.

## Discussion

In this study, we have identified a connectional axis of individual FC variability and further demonstrated its structural correlates and significance for development and cognition. Particularly, we found that the inter-individual FC variability is organized along a continuous axis at the connectional level across connectome edges. This connectional axis of FC variability progresses from within-network to between-network connections, and from connections within and between association networks to connections linking association and sensorimotor networks.

We found that this connectional variability axis aligns with structural connectivity variability across all connections. Moreover, we demonstrated that the connectional variability axis developed in youth with a lower axis slope. Besides, the slope of connectional variability axis was positively associated with individual differences in higher-order cognitive performance. These results were highly consistent between HCP-D and HCP-YA datasets and remained stable under a range of methodological variations, with further replication in an independent cohort of Chinese youth. Overall, we revealed a connectional axis of individual FC variability at the edge level, associated with structural connectome, refined in development, and implicated in cognition.

Prior studies have reported that individual variability in regional FC is heterogeneously distributed across the cortex along the sensorimotor-association cortical axis, with maximum variability in the association cortex and a gradual decrease along the axis towards the sensorimotor cortex (3, 8, 9). However, since regional FC is an aggregate measure, these studies lack the information provided by studying FC variability at the edge level. Our findings identified a connectional axis for the organization of edge-level individual FC variability across human connectome. This connectional axis defines a spectrum where FC variability decreases from within-network to between-network connections, and from connections within and between association networks to those between association and sensorimotor networks. Interestingly, we found that the lowest FC variability exists in the connections between association and sensorimotor networks rather than the connections within sensorimotor networks.

Our findings are consistent with the previous knowledge regarding the circuit architecture of the sensory and association cortices. Sensorimotor circuits primarily comprise circuits with canonical feedforward and feedback connectivity patterns (49). In contrast, higher-order association networks exhibit a noncanonical circuit architecture with distributed connections to many cortical areas (50), which may have contributed to the higher individual variability in the connections among these networks. One prominent theory suggests that the association connectivity may have emerged during evolution through the gradual untethering of connections from highly conserved patterns of sensorimotor connectivity and sensory-specific signaling molecules (51). This process may serve as the evolutionary basis of the FC variability axis from the edges connecting the sensorimotor network to the connections among the association networks. Moreover, in contrast to sensorimotor networks, association networks show much lower structural constraints in function (52), supporting the higher functional variations in the connections among association networks.

While previous studies identified the lowest variability within the sensorimotor cortex when examining regional FC variability, our findings reveal that the connections between association and sensorimotor networks have the lowest FC variability localizing at the bottom end of the connectional variability axis, rather than connections within the sensorimotor networks. The connections between sensorimotor and association networks are crucial for integrating sensory and motor inputs with higher-order cognitive processes (49, 53, 54), thus playing a fundamental role in survival by supporting essential functions such as perception and coordinated actions. Moreover, these connections typically involve long-range wiring compared to within-sensorimotor connections (55), indicating a higher metabolic and structural cost (56–58). Due to their essential role and higher cost, the brain may prioritize the stability and robustness of these sensorimotor-association connections (59), leading to lower inter-individual variability. Additionally, these integrative connections might be more evolutionarily conserved, as they are crucial for performing complex behaviors that require the synchronization of sensory, motor, and cognitive processes. Evolutionary pressure to maintain these connections may further contribute to their lower variability, ensuring reliable functionality across individuals.

We found that the pattern of connectional axis in FC variability was associated with that of structural communicability variability across the connectome edges. Previous studies have demonstrated that structural connectivity supports the functional dynamics, and the network communicability model explains functional propagation through structural connectivity using non-invasive neuroimaging and invasive direct electrical stimulation data (1, 6, 39, 40, 60), supporting the association between structural and functional variability. We observed that edges with higher individual FC variability also exhibited higher individual variability in structural communicability, localized in the connections among the higher-order association networks. Regions in the association cortex receive more diverse and polysynaptic inputs from different brain areas than sensorimotor regions (50, 51), which could result in higher individual variability in both structural connectivity and FC edges among association networks compared with edges connected to sensorimotor networks.

Our results provide a systematic examination of how the spatial patterning of edge-level FC variability is refined during youth, shedding light on broader neurodevelopmental processes. We observed that the spatial distribution of FC variability evolves with a notable decline in the slope of the connectional variability axis, driving by a pronounced reduction in FC variability of association connections, with relatively limited changes in connections involving sensorimotor areas. This finding aligns with the hierarchical cortical development framework, wherein youth is characterized by the maturation of higher-order association connections, while primary sensorimotor connections mature earlier (8, 25). Previous studies have highlighted a ubiquitous decline in short-range FC strength and an increase in long-range connectivity strength during youth (20, 61), both of which may contribute to the observed reduction in individual variability of FC among association networks. The decline in short-range FC strength likely reflects intense synaptic and axonal pruning procedure, which refines neural circuits by eliminating redundant or weaker connections (23), thereby reducing individual variability. Consistently, fMRI studies have shown a common developmental decline in connectivity strength among association networks at the group level across youth (19, 22). The long-range connection, which facilitate global information integration and support essential survival behaviors, may have been evolutionarily constrained to maintain stability. Therefore, the strengthening of long-range connectivity strength via myelination process (24) likely contributes to the reduction of FC variability. The synaptic and axonal pruning and myelination are both genetically encoded and shaped by experience-dependent activity (23, 24). While distinct individual experience can increase variability, the common genetically encoded processes and shared environmental exposures—such social interactions and learning—reduce variability across individuals. The refinement of short- and long-range connectivity promotes greater global integration and local segregation of higher-order association networks, a general developmental principle consistently observed during youth (21). Together, these developmental findings on the spatial patterning of individual FC variability integrate well with existing neurodevelopmental frameworks, providing a deeper understanding of the distinct trajectories of connectivity maturation across the human connectome.

Finally, we found that the slope of the connectional variability axis was positively correlated with higher-order cognition, a composite score of multiple cognitive tasks (47), such as working memory, inhibition, and flexibility. We further demonstrated that better performance in higher-order cognition is associated with enhanced among-association variabilities at the top end of the connectional variability axis and a decline in association-sensorimotor variabilities at the bottom end. The higher-order cognitive abilities mainly rely on the association network, which is composed of a spatially distributed set of brain regions that span the frontal, parietal, and temporal areas (25, 62–64). Prior studies have proposed that, along with the emergence of higher-order cognition, association networks untethered from rigid canonical circuits have developed a highly variable and distributed pattern of connections during evolution (51). From this evolutionary perspective, higher variability in the connections among association networks, including fronto-parietal, default mode and attention networks, could explain improved performance in higher-order cognition. On the other hand, individuals with better higher-order cognition are likely to exhibit lower variability in connections with sensorimotor regions. This finding could explain how sensorimotor connections maintain stability, supporting the primary sensory and motor functions critical to survival and ensuring robustness against vulnerability.

Several methodological considerations must be considered when interpreting these findings. First, our analyses focused on the connections between cortical regions. Future studies should evaluate the variability axis of edges connected to subcortical and cerebellar structures. Second, accurately reconstructing individuals’ white matter structural connectivity poses significant challenges. Previous studies have shown that dMRI-based fiber tractography can yield both false positives and false negatives (65). In this study, we employed advanced probabilistic fiber tractography utilizing multi-shell, multi-tissue constrained spherical deconvolution, alongside anatomically constrained tractography (43) and spherical deconvolution-informed filtering of tractograms (44) to enhance biological accuracy (66). Third, while the correlation between FC variability and SC variability is relatively weak, our results are consistent with the effect sizes reported in prior literature evaluating the similarity between cortical maps or connectome matrices (17, 18).

These limitations notwithstanding, we identified a stable and reproducible connectional axis of edge-level individual FC variability across connectome edges. This connectional axis of FC variability aligned with spatial variation of structural connectivity variability. While the human connectome exhibits extreme complexity, the connectional axis captures an important organizational motif, which might be fundamental for the human connectome. Moreover, the connectional axis of FC variability was refined during development and was related to higher-order cognition, therefore this variability axis may be important for understanding the neurodevelopmental substrates of mental disorders, as well as in the design of personalized targets for neuromodulatory interventions.

## Materials and Methods

Methods are summarized below; see SI Appendix for further detail.

### Study cohorts and participants

This study utilized multi-modal neuroimaging data, including T1-weighted structural MRI, resting-state functional MRI (fMRI), and diffusion MRI, from 339 unrelated participants (183 females, aged 22–37) in the HCP-YA dataset (Release S900) (32) and 633 participants (339 females, aged 8–21) obtained from the HCP-D dataset (Release 2.0) (33). The HCP-YA and HCP-D studies were approved by the Washington University Institutional Review Board, and informed consent was obtained from all participants. After data quality control, 275 participants from HCP-YA and 486 participants from HCP-D were included in subsequent analyses. Four resting-state fMRI runs of each individual from HCP-YA and four resting-state fMRI runs of each individual from HCP-D were used to estimate intra- and inter-individual variability.

### FC measured using BOLD fMRI data

Minimally preprocessed T1-weighted structural and functional MRI data were acquired from the HCP-D and HCP-YA datasets (67). Preprocessed fMRI data were then post-processed using the extensible connectivity pipelines (XCP-D, 0.2.2) (68). The regional BOLD time series were extracted using an a priori Schaefer parcellation with 400 parcels (34). FC was calculated as the Pearson correlation coefficient between each pair of regional time series, followed by Fisher’s r-to-z transformation, resulting in a 400×400 FC matrix for each participant. The parcels were then mapped to six large-scale functional networks from the Yeo atlas (37), excluding the limbic network, as previous studies have consistently reported substantial signal loss in this network(38). This resulted in a 374×374 FC matrix with 69,751 unique edges per participant.

### Inter-individual variability of FC and the connectional variability axis

We estimated inter-individual variability in FC edges, and a split-session approach (69) was adopted to estimate intra-individual variability. For the HCP-YA dataset, we split four resting-state fMRI runs into twelve sessions, producing twelve FC matrices per participant. In the HCP-D dataset, we concatenated four resting-state and split them into eight sessions, yielding eight FC matrices per participant.

We calculated inter- and intra-individual variability of each FC edge using a linear mixed-effects model with the R package Rex (35). This approach produced two 374×374 variability matrices for inter- and intra-individual variability for each dataset. Subsequently, we used the intra-class correlation to measure the adjusted inter-individual variability while accounting for intra-individual variability (3).

To simplify, we averaged FC variability within and between the six large-scale networks, creating a 6×6 inter-individual variability matrix at the network level, represented by 21 unique FC variability values. We termed the axis in the individual variability matrix, which progresses from connections within and between association networks to the connections between association and sensorimotor networks, the “connectional axis” of FC variability. We further ranked the 21 FC variability values from highest to lowest, and calculated the slope of the connectional variability axis using linear regression to measure the heterogeneity of connectional variability across the connectome (See Supplementary Methods for details).

### Structural connectome construction with diffusion MRI based white matter tractography

Diffusion MRI data were preprocessed using Mrtrix3 (70) and QSIPrep (71). Whole-brain white matter tracts were reconstructed to create structural connectivity (SC) using the Schaefer-400 atlas. Structural edge weights, representing the number of streamlines between regions, were normalized by dividing by the average volume of the two regions (72) and then log-transformed. A consistency-based thresholding method was applied to reduce spurious connections (73), using the 75th percentile for edge weight coefficient of variation (52). To account for both direct and indirect pathways, we calculated the communicability of the SC matrix for each participant (40, 41). The mean absolute deviation of structural communicability across participants for each edge was used to define inter-individual variability in structural communicability.

### Development of connectional variability axis in youth

To explore the maturation of connectional variability in youth, we divided HCP-D participants (aged 8–21) into subgroups and calculated the inter-individual variability matrix for each group. Using a sliding window approach (74), 486 participants were sorted by age and divided into 88 overlapping groups (window length = 50, step size = 5), with average ages ranging from 9.2 to 21.2 (**Fig. 3a**). We then calculated the FC variability matrix for each group.

We estimated the “connectional axis slope” of FC variability for each of the 88 groups and evaluated developmental changes in the axis slope during youth. To assess both linear and nonlinear effects of development, we used a generalized additive model (GAM), with connectional axis slope as the dependent variable and age as a smooth term, controlling for sex and motion.

The significance of the age term was assessed through analysis of variance (ANOVA) that compared the full GAM model to a nested, reduced model with no age term (75). The overall effect was measured by the partial *R*^2^ between the full GAM and reduced models (effect magnitude), and we signed the partial *R*^2^ by the sign of the age coefficient from an equivalent linear model (effect direction) (22, 75). Finally, to determine which connections drive the change in axis slope, we constructed the same GAM model for each network-level FC and estimated the developmental effects of its FC variability.

### Association between connectional variability axis and cognitive performance

We examined how the connectional variability axis associates with cognitive performances in the HCP-D and HCP-YA datasets. We used the composite score of fluid cognition from the NIH Toolbox Cognition Battery to quantify the participants’ cognitive abilities (47). This analysis included 352 participants from the HCP-D and 274 participants from the HCP-YA with fluid cognition composite scores.

We ranked the participants based on their cognitive scores and then employed a sliding window approach (window length = 50, step size = 5) to divide them into multiple groups with increasing cognitive scores for each dataset (**Fig. 4a**). In total, 61 and 45 groups were obtained for HCP-D and HCP-YA, respectively. We estimated the inter-individual FC variability in each group and calculated the connectional axis slope with network-level average variability. A GAM analysis was performed to evaluate the relationship between the connectional axis slope of FC variability and fluid cognition composite scores while controlling for the linear and nonlinear effects of age, as well as the effects of sex and in-scanner motion.

The *t* value associated with the cognition term in the GAM represents the direction of the slope-cognition association, and its significance (*P* value) was also derived from the GAM model. The partial *R*^2^ between the full GAM and reduced models with no cognition term was calculated as the effect size. Similarly, to determine which connections drive the change in axis slope, we constructed the same GAM model for each network-level FC and estimated the cognitive effects of its FC variability.

### Replication in an independent Chinese youth dataset

We further included an independent Chinese youth cohort, the Youth Executive function and Neurodevelopment (YEN) study (*n* = 253, 123 females, aged 6–22), to replicate all primary findings. The YEN study was approved by the Human Research Ethics Committee of the Chinese Institute for Brain Research, Beijing (CIBR-HREC-20230317-001). Written assent and informed consent were obtained from all participants. For participants under 18 years of age, consent was also obtained from their parents or legal guardians. After data quality control, 218 participants were included in subsequent analyses. We conducted all primary analyses on this dataset the same procedures as applied to the HCP-D dataset. Supplementary Materials for details of data parameters, preprocessing steps, and analysis procedures.

### Data, Materials, and Software Availability

All code used to perform the analyses in this study, functional connectivity, individual variability in functional connectivity, and individual variability in structural communicability matrices can be found at https://github.com/CuiLabCIBR/Connectional_Variability_Axis. The HCP-YA and HCP-D datasets are available at https://db.humanconnectome.org/.

## Acknowledgments

This work is supported by the STI 2030-Major Projects (2022ZD0211300), CAMS Innovation Fund for Medical Sciences (2024-I2M-ZD-013), Non-profit Central Research Institute Fund of Chinese Academy of Medical Sciences (2024-RC416-02), Beijing Nova Program (Z211100002121002), and CIBR funds to Z.C.. V.J.S. is supported by the National Institute of Mental Health, USA (grant number T32MH016804). Data were provided [in part] by the Human Connectome Project, WU-Minn Consortium (Principal Investigators: David Van Essen and Kamil Ugurbil; 1U54MH091657) funded by the 16 NIH Institutes and Centers that support the NIH Blueprint for Neuroscience Research; and by the McDonnell Center for Systems Neuroscience at Washington University. The HCP-Development 2.0 Release data used in this report came from DOI: 10.15154/1520708. Research reported in this publication was supported by the National Institute of Mental Health of the National Institutes of Health under Award Number U01MH109589 and by funds provided by the McDonnell Center for Systems Neuroscience at Washington University in St. Louis. The Youth Executive function and Neurodevelopment data were provided by our lab in CIBR, supported by the STI 2030-Major Projects (2022ZD0211300).

