## Supporting Information for "Connectional axis of individual functional variability: Patterns, structural correlates, and relevance for development and cognition"

**This PDF file includes:**

SI Methods
SI Results
Figures S1 to S3
Tables S1 to S3
SI References

### SI Methods

#### Study cohorts and participants

**HCP young adult (HCP-YA) dataset**. This study utilized multi-modal neuroimaging data, including T1-weighted structural MRI, resting-state functional MRI (fMRI), and diffusion MRI, from 339 unrelated participants (183 females, aged 22–37) in the HCP-YA dataset (Release S900) (1). The HCP-YA study was approved by the Washington University Institutional Review Board, and informed consent was obtained from all participants. All imaging data were acquired on a Siemens 3T Skyra scanner. Structural MRI data were scanned using a resolution of 0.7mm isotropic resolution. Two resting-state fMRI sessions with two runs in each session, left-right (LR) and right-left (RL) phase-encodings, were acquired for each participant with a 2 mm isotropic resolution, TR = 0.72 s. Each resting-state run included 1,200 frames and was approximately 15 min long. Diffusion MRI data were acquired in two runs with opposite phase-encoding directions for each participant. Each run included 270 noncollinear directions with three nonzero shells (b = 1,000, 2,000, and 3,000 s/mm$^2$). Further details regarding the HCP-YA dataset and MRI acquisition parameters have been described in prior study (1).

**HCP development (HCP-D) dataset**. This study also included 633 participants (339 females, aged 8–21) obtained from the HCP-D dataset (Release 2.0) (2), including T1-weighted structural MRI, resting-state fMRI, and diffusion MRI. The HCP-D study was approved by the institutional review boards at Washington University (IRB #201603135). Written informed consent and assent were obtained from participants over 18 years of age and parents of participants under 18 years. All data were collected on a 3T Siemens Prisma scanner. The resolution for structural MRI data was 0.8 mm isotropic. Two resting-state fMRI sessions were acquired for each participant, with two runs in each session, using anterior-posterior (AP) and posterior-anterior (PA) phase encoding. Each resting-state run lasted approximately 6.5 min with 488 frames, TR = 0.8 s. The diffusion MRI data included two sessions, each with two shells (b = 1500, 3000 s/mm$^2$) and 185 diffusion-weighted directions. Further details about the HCP-D dataset have been described in a previous study (3).

**Replication dataset: Youth Executive function and Neurodevelopment (YEN) study**. The YEN is an ongoing study collecting multi-modal neuroimaging and phenotype data from Chinese children and adolescents. This study included 253 participants (123 females, aged 6–22) who underwent T1-weighted structural MRI, resting-state fMRI, and diffusion MRI scans. The YEN study was approved by the Human Research Ethics Committee of the Chinese Institute for Brain Research, Beijing (CIBR-HREC-20230317-001). Written assent and informed consent were obtained from all participants. For participants under 18 years of age, consent was also obtained from their parents or legal guardians. All MRI data were acquired using a 3T Siemens Prisma scanner. Structural MRI scans were obtained with a 1.0 mm isotropic resolution. Each participant completed three resting-state fMRI runs (TR = 2 s), each lasting 6 minutes, accompanied by a spin-echo field map. Additionally, diffusion MRI data were acquired with 120 gradient directions (b-values: 500, 1000, 2000, and 3000 s/mm$^2$) along with a reverse phase-encoded b0 image for distortion correction.

#### Structural and functional MRI data processing

HCP-YA and HCP-D preprocessing
Minimally preprocessed T1-weighted structural and functional MRI data were acquired from the HCP-YA and HCP-D datasets (4). Structural data were corrected for intensity non-uniformity, skull-stripped, and used to reconstruct the cortical surface. Volume-based structural images were segmented into cerebrospinal fluid (CSF), white matter, and gray matter, and then spatially normalized to the standard Montreal Neurological Institute (MNI) space. The fMRI data were preprocessed with slice-timing correction, motion correction, distortion correction, co-registration to structural data, normalization to the MNI space, and projection to the cortical surface.

Functional time series were resampled to FreeSurfer's fsaverage space, and grayordinates files containing 91k samples were generated.

YEN preprocessing
T1-weighted structural and functional MRI data were preprocessed using the robust pipeline in fMRIPrep version 20.2.0 (https://fmriprep.org/en/20.2.0/) (5). For T1-weighted images, preprocessing included intensity normalization, skull stripping, brain tissue segmentation, spatial normalization, and surface reconstruction using FreeSurfer v6.0.1 (https://surfer.nmr.mgh.harvard.edu/fswiki) (6). Then, fMRI data underwent motion correction, distortion correction, co-registration to the T1-weighted data, normalization, and projection to the cortical surface. The processed functional time series were resampled to FreeSurfer's fsaverage space and the 32k fsLR space.

Preprocessed fMRI data were then post-processed using the extensible connectivity pipelines (XCP-D 0.2.2; https://xcp-d.readthedocs.io/en/0.2.2/) (7). We regressed 36 nuisance regressors out from the BOLD data, including six motion parameters, global signal, mean white matter signal, and mean CSF signal, along with their temporal derivatives, quadratic terms, and quadratic derivatives (8). Residual timeseries were then band-pass filtered (0.01–0.08 Hz).

We excluded participants using two criteria to minimize potential head motion effects (9). First, an fMRI run was ruled out when over 25% of the frames had an FD > 0.3 mm. Second, for each fMRI run, we calculated the mean FD distribution by pooling frames from all participants and derived the third quartile (Q3) and interquartile range (IQR) of this distribution. Runs with mean FD greater than "Q3+1.5×IQR" were excluded. We only included participants whose fMRI runs satisfied these criteria in HCP-YA (four resting-state fMRI runs), HCP-D (four resting-state fMRI runs), and YEN (three resting-state fMRI runs) datasets. We excluded 64 HCP-YA, 139 HCP-D, and 35 YEN participants based on these two criteria. Additionally, 8 participants from the HCP-D were excluded because of incomplete fMRI runs (less than 4 runs). Consequently, 275 participants (146 females, aged 22–35) from the HCP-YA, 486 participants (265 females, aged 8–21) from the HCP-D, and 218 participants (113 females, aged 6–22) from the YEN datasets were included in subsequent analyses.

**Diffusion MRI data processing**

Diffusion MRI data from the HCP-D and YEN datasets were preprocessed using QSIPrep (https://qsiprep.readthedocs.io/), an integrative platform for preprocessing and reconstructing diffusion MRI data (10), which includes tools from Mrtrix3 (https://www.mrtrix.org/) (11). Prior to preprocessing, we concatenated the EPI runs with opposing phase-encoded directions and extracted the frames where the b-value was < 100 s/mm$^2$ as the b0 image. Next, we applied MP-PCA denoising, Gibbs unringing, and B1 field inhomogeneity correction using the Mrtrix3's *dwidenoise* (12), *mrdegibbs* (13), and *dwibiascorrect* (14) functions, respectively. An FSL eddy was used for head motion and eddy current corrections (15). Finally, the preprocessed diffusion-weighted imaging (DWI) data were resampled to the ACPC space at an isotropic resolution of 1.5 mm. We used minimally preprocessed diffusion MRI data from the HCP-YA dataset. The minimal preprocessing pipeline included b0 image intensity normalization across runs, EPI distortion correction, eddy current and motion correction, gradient nonlinearity correction, and registration to the native structural space (1.25 mm). The processed diffusion MRI data were further corrected for B1 field inhomogeneity using Mrtrix3. Particularly, a total of 8 participants from the HCP-D dataset and 6 participants from the YEN dataset were excluded from subsequent structural connectivity analyses due to incomplete DWI data, failures in diffusion data preprocessing or the identification of isolated regions after tractography.

**FC measured using BOLD fMRI data**

First, the regional BOLD time series was extracted using an a priori Schaefer parcellation with 400 parcels (16). FC was calculated as the Pearson correlation coefficient between each pair of regional timeseries, resulting in a 400×400 symmetrical FC matrix for each participant. Fisher's z-transformation was applied to each FC value in the matrix. Subsequently, the parcels were mapped onto seven canonical large-scale functional networks from the Yeo atlas (17). We excluded the limbic network in the following analyses, as previous studies have consistently reported substantial signal loss in this network (18, 19), especially in the orbitofrontal and ventral temporal cortices. Our analysis contained 374 parcels from six functional networks, including the visual (VS), somatomotor (SM), dorsal attention (DA), ventral attention (VA), frontoparietal (FP), and default mode (DM) networks. Finally, a 374×374 symmetrical FC matrix with 69,751 unique edges was obtained for each participant. The Schafer atlas with 374 cortical regions was also used to construct connectomes from other data modalities.

**Inter-individual variability of FC and the connectional variability axis**

We estimated inter-individual variability in FC edges and evaluated the organizing principle at the connectional edge level. Due to the limited number of scans per participant, a split-session approach (20) was adopted to estimate intra-individual variability. For the HCP-YA dataset, we split each of the four resting-state fMRI runs into three sessions with equal-length time series, producing twelve sessions per participant and acquiring twelve FC matrices for each participant. We concatenated four resting-state fMRI runs for the HCP-D dataset and split them into eight sessions. We acquired eight FC matrices for each participant from the HCP-D dataset.

Inter- and intra-individual variability of each FC edge (69,751 unique edges in total) were calculated using a linear mixed-effects (LME) model for each dataset (21). The LME model was implemented via the R package Rex (https://github.com/TingsterX/Reliability_Explorer), which was designed to facilitate the robust measurements of individual variations (21). This model captures fixed and random effects, assuming that the observed response variable and residual term follow a normal distribution with zero mean and a specific variance. Previous studies have used this approach to measure inter- and intra-individual variability (22, 23). Specifically, for each FC edge, the LME model is expressed as:

$$FC_{it} = \mu_0 + \lambda_i + \alpha_t + \epsilon_{it}, \text{ where } \lambda_i \sim N(0, \sigma_\lambda^2),\ \epsilon_t \sim N(0, \sigma_\epsilon^2). \quad (1)$$

Here, *i* identifies the individual, *t* indicates the session, $\lambda_i$ and $\alpha_t$ measures random effect and fixed effect, respectively, and $\epsilon_{it}$ is the residual term. The observed individual variation $\sigma_{FC}^2$ between $FC_{it}$ can be decomposed into real inter-individual variation $\sigma_b^2$ across participants and the intra-individual variations $\sigma_w^2$ across sessions (captured by the residual variation $\sigma_\epsilon^2$). $\sigma_b^2$ and $\sigma_w^2$ represent inter- and intra-individual variabilities for one specific FC edge, which were used in our following analyses.

We obtained two 374×374 variability matrices for inter- and intra-individual variability for each dataset. Subsequently, we used the intra-class correlation (ICC, $\frac{\sigma_b^2}{\sigma_b^2+\sigma_w^2}$) to measure the adjusted inter-individual variability while accounting for intra-individual variability (24). Thus, we acquired a matrix of inter-individual variability for the HCP-D and HCP-YA datasets. Next, we compared the similarity in the connectional axis between the inter-individual variability matrices from the HCP-D dataset and HCP-YA datasets using Spearman's rank correlation.

We termed the axis in the individual variability matrix, which progresses from connections within and between association networks to the connections between association and sensorimotor networks, the "connectional axis" of FC variability. For a more straightforward illustration, we averaged the FC variability within the six large-scale networks and between all network pairs, yielding a 6×6 symmetrical inter-individual FC variability matrix at the network level. The connectional axis of this variability matrix can be uniquely represented by six within-network FC variability values (diagonal elements) and 15 between-network FC variability values (nondiagonal elements). Twenty-one unique FC variability values were defined as the profile of the connectional variability axis. We calculated Spearman's rank correlation between the variability axis profiles from the HCP-D and HCP-YA datasets. To visualize the connectional axis of FC variability, we sorted the within-network and between-network FC variability values in descending order for each functional network separately. We further ranked the 21 FC variability values from highest to lowest, and calculated the slope of the connectional variability axis using linear regression to measure the heterogeneity of connectional variability across the connectome. Specifically, the linear model was constructed as $Y = \beta_0 + \beta_1 * X$, where Y represents the 21 ranked FC variability values, and the X is the rank ranging from 1 to 21 (scaled from 0 to 1 by dividing each rank by 21), $\beta_0$ is the intercept, and $\beta_1$ measures the slope of the connectional variability axis. Finally, we grouped the 69,751 unique edges by connection type (within-sensorimotor, within-association, and between sensorimotor-association) and compared their differences in FC variability using a permutation test (10,000 iterations).

**Structural connectome construction with diffusion MRI based white matter tractography**

Based on the preprocessed diffusion MRI data, a connectome of structural connectivity (SC) was generated for each participant using reconstructed whole-brain whiter matter tracts. Reconstruction was done using MRtrix3, which implements a multi-shell, multitissue constrained spherical deconvolution (CSD) to estimate the fiber orientation distribution (FOD) for each voxel (25). We used an anatomically constrained tractography (ACT) framework to improve the biological accuracy of fiber reconstruction (26). Specifically, tractography was performed using *tckgen*, which generates 10 million streamlines (length ranging from 30 to 250 mm, FOD power = 0.33) using improved probabilistic streamlines tractography (iFOD2) based on the second-order integration over FOD. The ACT approach integrates anatomical constraints to guide the tractography processes by using the tissue segmentation (e.g., gray matter, white matter, and cerebrospinal fluid) to ensure the fibers are confined within anatomically plausible regions (26). We selectively filtered the streamlines from the tractogram based on the spherical deconvolution of the diffusion signal and estimated the streamline weights using the command *tcksift2* (27). Next, a structural connectivity matrix was constructed using *tck2connectome* with the Schaefer-400 atlas. A radial search was performed from each streamline endpoint to identify the nearest gray matter node, with a maximum search radius of 2 mm (27).

We visualized the reconstructed fiber tracts on translucent cortical pial surfaces for four randomly selected participants—two from the HCP-D dataset and two from the HCP-YA dataset (**Fig. S1**). Given that the probabilistic tractography generated 10 million streamlines, which were too dense for effective visualization, 10,000 streamlines were randomly selected using the MRtrix3 command *tckedit* and then visualized using 3D slicer (https://www.slicer.org/). The visualizations demonstrate that the reconstructed white matter tracts align with known anatomical structures, including the corona radiata, posterior limbs of the internal capsule, inferior longitudinal fasciculus, and corpus callosum.

The weight of each structural connectivity indicates the number of streamlines connecting two regions. Edge weights were normalized by dividing the average volume of the two regions (28), and then log-transformed. Specifically, we employed a consistency-based thresholding method (29) to mitigate the risk of spurious white matter connections due to probabilistic tractography. In accordance with previous studies (30, 31), we first computed the coefficient of variation (CV) for edge weight across participants, and subsequently, we applied a threshold to individual structural connectivity matrices at the 75th percentile for edge weight CV.

Considering that the white matter structural connectome accounts only for direct functional communication, we calculated the communicability of the structural connectivity matrix for each participant to include both the direct and indirect pathways (32, 33). Communicability quantifies the capacity of two regions to communicate through other regions by pathways of all possible topological lengths. Specifically, let *A* denotes the weighted adjacency matrix of the individual structural connectome. The communicability between two regions *i* and *j* is calculated as $CMY_{ij} = (\exp(D^{-1/2}AD^{-1/2}))_{ij}$, where $D = diag(\sum_{k=1}^{N} a_{ik})$ is a diagonal matrix in which the diagonal elements represent the strength of each node. We calculated the mean absolute deviation of structural communicability across all participants for each edge and defined it as the inter-individual variability of structural communicability across all participants. We evaluated the alignment between FC variability and structural communicability variability across all edges by computing Spearman's rank correlation of the connection strength between the communicability variability and FC variability. We employed a bootstrap approach (1,000 repetitions) to estimate the 95% confidence interval (CI) of Spearman's rank correlation using the R package bootcorci (https://github.com/GRousselet/bootcorci) (34).

**Null models for between-connectome correlations**

To evaluate the significance of the correlation between the two connectomes across edges, we performed permutation testing by generating null connectomes, as in the spatial permutation test, to compare cortical properties (35). The spin test was designed to assess the spatial similarity between two cortical maps while considering autocorrelations in the brain data. This is achieved by projecting the brain surface onto a spherical space, rotating the sphere, and projecting it back onto the surface (36). Here, we utilized the Schaefer atlas, comprising 400 cortical regions with corresponding node IDs ranging from 1 to 400 (i.e., [1, 2, ..., 400]). To create the null connectomes, we shuffled the 400 cortical regions by rotating the Schaefer atlas. We then reordered the rows and columns of the original matrix based on the shuffled node IDs (e.g., [134, 25, ..., 365]). This procedure preserved the mean and variance of the matrix while ensuring that the regional profile (i.e., the matrix column, 400×1) included all 400 cortical regions. Repeating this process, we constructed a null distribution by generating 10,000 randomized individual variability connectomes. Next, we calculated Spearman's rank correlations between the randomized FC variability and the structural communicability variability. We compared the correlation values obtained from the empirical individual variability matrix with those acquired using the null connectomes to determine the significance level, and the $P$ value of this permutation testing ($P_{perm}$) was reported.

**Development of connectional variability axis in youth**

To explore the maturation of the connectional variability axis in youth, we divided the HCP-D participants (aged 8–21 years old) into multiple subgroups and calculated the inter-individual variability matrix for each group. We balanced the number of groups and group sizes using a sliding window approach (37) to ensure sufficient statistical power and accurate estimation of individual variability. Specifically, 486 participants from the HCP-D dataset were sorted in ascending order by age and then divided into overlapping groups (**Fig. 3a**). We set the window length to 50 participants with a step size of 5 participants. We obtained 88 groups, with the average age ranging from 9.2 to 21.2. We then calculated the inter-individual variability across all participants and acquired a 374×374 FC variability matrix for each group. A group average age was assigned for each group. The sex of each group was computed as the percentage of males. Head motion was calculated by averaging mean FD across all individuals and all runs within each group.

We evaluated how the connectional axis of individual FC variability developed in youth. The connectional variability axis represents the ranking of the 21 within-network and between-network average variabilities. The peak of this axis compromised connections within- and between-association networks, whereas the between-network edges linking the sensorimotor networks were at the base of the axis. We estimated the "connectional axis slope" of FC variability for each of the 88 groups and evaluated developmental changes in the axis slope during youth. To model both linear and nonlinear developmental effects of connectional axis slope, we used a generalized additive model (GAM) (38) with penalized splines, which estimates nonlinearities using restricted maximum likelihood (REML) and penalizes nonlinearity to avoid overfitting data. GAM was fit with the connectional axis slope was modeled as the dependent variable. The group age was modeled as a smooth term while group sex and group in-scanner head motion as model covariates as follows:

$$Connectional\ axis\ slope \sim s(Age,\ k = 3) + Sex + Motion \quad (2)$$

where *s*() is the spline basis function, and *k* determines the maximum basis complexity. We set *k* to 3 because recent studies have suggested that *k* = 3 can capture the nonlinear effects of development while preventing overfitting (39, 40). The significance of the association between the connectional axis slope of FC variability and age was assessed through analysis of variance (ANOVA) that compared the full GAM model to a nested, reduced model with no age term (40). A significant result indicated that the residual deviance was significantly lower when a smooth term for age was included in the model, as assessed using the chi-squared test statistic. To determine the overall magnitude and direction of the association between the connectional axis slope of FC variability and age, we calculated the partial $R^2$ between the full GAM and reduced models (effect magnitude), and signed the partial $R^2$ by the sign of the age coefficient from an equivalent linear model (effect direction) (40, 41). Finally, to determine which connections drive the change in axis slope, we constructed the same GAM model for each network-level FC and estimated the developmental effects of its FC variability.

**Association between connectional variability axis and cognitive performance**

We evaluated the association between the FC variability axis and individual differences in cognition. Specifically, we examined how the connectional axis slope of FC variability varied among participants with different cognitive performances in the HCP-D and HCP-YA datasets. We used the composite score of fluid cognition from the NIH Toolbox Cognition Battery to quantify the participants' cognitive abilities (42). The fluid cognition composite score was obtained by averaging the normalized scores from multiple cognitive tasks, including flanker inhibition, dimensional change card sort (flexibility), picture sequence memory, list sorting working memory, and pattern comparison. This analysis included 352 participants from the HCP-D and 274 participants from the HCP-YA with fluid cognition composite scores.

We ranked the participants based on their cognitive scores and then employed a sliding window approach (window length = 50, step size = 5) to divide them into multiple groups with increasing cognitive scores (**Fig. 4a**). In total, 61 and 45 groups were obtained for the HCP-D dataset and the HCP-YA dataset, respectively. We estimated the inter-individual FC variability in each group and calculated the connectional axis slope with network-level average variability. A GAM analysis was performed to evaluate the relationship between the connectional axis slope of FC variability and fluid cognition composite scores while controlling for the linear and nonlinear effects of age, as well as the effects of sex and in-scanner motion. The equation for GAM is as follows:

$$Connectional\ axis\ slope \sim Cognition + s(Age,\ k = 3) + Sex + Motion \quad (3)$$

The *t* value associated with the cognition term in the GAM represents the direction of the slope-cognition association, and its significance (*P* value) was also derived from the GAM model. The partial $R^2$ between the full GAM and reduced models with no cognition term was calculated as the effect size. Similarly, to determine which connections drive the change in axis slope, we constructed the same GAM model for each network-level FC and estimated the cognitive effects of its FC variability.

**Sensitivity analyses**

FC strength can influence the scale of individual FC variability, wherein edges with stronger FC strength may exhibit larger individual FC variability. Similarly, the strength of structural communicability may also influence the association between FC variability and communicability variability. Therefore, we controlled for both mean FC and communicability strength, and re-estimated the spatial alignment between the FC variability matrix and SC variability matrix across edges. This procedure was performed for both the HCP-D and HCP-YA datasets. Finally, in the analyses related to development and cognition, we regressed out the mean FC strength from the variability matrix for each group obtained by the sliding window approach, and re-evaluated effects of development and cognition on the connectional variability axis.

Previous studies have reported that adjacent brain regions tend to be associated with analogous molecular and structural bases and exhibit similar functions (43-45). To account for the effects of interregional distance, we calculated the Euclidean distance between every two cortical regions in the MNI space for the Schaefer-400 atlas. We regressed out the interregional distance from the FC variability matrix and SC variability matrix and re-estimated their correlations. Similarly, we regressed out the interregional distance from the individual variability matrices of each group before evaluating the developmental and cognitive effects of the connectional variability axis.

We also replicated our main findings using the HCP's multi-modal parcellation (46), a widely acknowledged brain parcellation. Additionally, as the global signal shows network-specific effects and changes across development (47, 48), we evaluated the sensitivity of our findings when global signal regression was not applied.

To further minimize the potential influence of spurious structural connections on the association between individual FC variability and structural communicability variability, we applied a minimum threshold of streamline density within participants, retaining only streamlines with the largest weights. We tested a range of streamline density thresholds from 5% to 25% in increments of 5%. For example, at a density threshold of 5%, connections with streamline counts in the top 5% were set to 1, and all others were set to 0 for each individual independently. Communicability was then computed based on the binary SC network for each participant, and structural communicability variability was assessed across individuals.

Finally, we confirmed that our developmental and cognitive analyses were robust to the choice of window length and step size in the sliding window configurations. Specifically, we tested six combinations comprising three window lengths (40, 50, and 60 participants) and two step sizes (5 and 10 participants).

**Replication in an independent Chinese youth dataset**

We replicated all primary findings using an independent Chinese youth dataset from the YEN study. For each individual, three resting-state fMRI runs without splitting were used to compute three FC matrices, as there are only 180 frames for each run. These FC matrices were then utilized to estimate adjusted inter-individual FC variability, accounting for intra-individual variability, within the YEN dataset. The individual variability of structural communicability was also estimated in the YEN dataset and correlated with individual FC variability using Spearman's rank correlation.

For the developmental analyses, with the same sliding window approach and parameters (window length = 50, step size = 5), the 218 participants were divided into 34 groups and individual FC variability was estimated within each group. Then, the same GAM model used for the HCP-D dataset was employed to assess the relationship between the connectional axis slope of FC variability and age.

Finally, we examined the association between the connectional variability axis and higher-order cognition. The YEN study evaluated executive function across three components—working memory, inhibitory control, and cognitive flexibility—using eight tasks. Task performance metrics included accuracy rates for the working memory tasks (spatial 2-back, number 2-back, keep track), stop-signal response time for the stop signal task (based on (49)), and a 2-vector score integrating accuracy and reaction time for inhibitory control tasks (flanker, color Stroop) and cognitive flexibility tasks (dots triangles, dimensional change card sort) (50). A total of 179 participants with complete behavioral scores across all assessments were included in the analyses. A composite executive function score was derived by averaging z-scores for individual task performances, followed by normalization to a mean of 100 and a standard deviation of 15

(50). The 179 participants were divided into 26 groups using the same sliding window approach, and individual FC variability was estimated within each group. Subsequently, the same GAM model utilized in previous cognitive analyses was used to evaluate the relationship between the connectional axis slope of FC variability and executive functions.

### SI Results

#### Sensitivity analyses

We conducted sensitivity analyses to evaluate the robustness of our findings. We first evaluated the influence of FC strength in our findings by regressing out FC strength from the individual FC variability and subsequently repeated all the analyses. For the analysis of structural communicability variability, we additionally regressed out the structural communicability strength from the structural communicability variability and FC variability. We found all the results remain significant after controlling for FC strength. As shown in **Fig. S2a**,**b**, we found that the pattern of individual FC variability remained significantly correlated with structural communicability variability across connectome edges (HCP-D: rho = 0.17, $P_{perm} < 0.0001$; HCP-YA: rho = 0.21, $P_{perm} < 0.0001$). Additionally, after regressing out the FC strength from the individual FC variability of each of the 88 age groups, we observed a significant decline in the connectional axis slope during youth (partial $R^2 = -0.08$, $P = 0.008$, **Fig. S3a**). Similarly, the connectional axis slope of FC variability remained significantly correlated with the fluid cognition in both the HCP-D ($t = 2.40$, $P = 0.02$, partial $R^2 = 0.09$, **Fig. S3b**) and HCP-YA ($t = 8.90$, $P = 6.04\times10^{-11}$, partial $R^2 = 0.68$, **Fig. S3c**) datasets.

Previous studies have described the influence of physical distance between regions on structural connectivity and FC (45, 51). To minimize the effect of spatial proximity, we regressed out the Euclidean distance between regions from the individual variability matrix and structural communicability variability. As shown in **Fig. S2c,d**, the pattern of individual FC variability remained significantly correlated with structural communicability variability across connectome edges (HCP-D: rho = 0.31, $P_{perm} < 0.0001$; HCP-YA: rho = 0.26, $P_{perm} < 0.0001$). Consistently, we observed that the connectional axis slope of FC variability was significantly associated with age in youth (partial $R^2 = -0.32$, $P = 3.66\times10^{-10}$, **Fig. S3d**). The connectional axis slope of FC variability remained significantly associated with the fluid cognition composite score in both the HCP-D ($t = 3.29$, $P = 0.002$, partial $R^2 = 0.14$, **Fig. S3e**) and HCP-YA ($t = 5.95$, $P = 5.94\times10^{-7}$, partial $R^2 = 0.48$, **Fig. S3f**) datasets.

Furthermore, we verified that our main findings were not sensitive to brain parcellation. To this end, we replicated our main findings using the HCP's multi-modal parcellation (46), a widely acknowledged brain parcellation consisting of 360 cortical regions (Glasser-360). As depicted in **Fig. S2e,f**, our main findings remained consistent and statistically significant. For instance, the FC variability was still positively correlated with the structural communicability variability in both the HCP-D (rho = 0.25, $P_{perm} < 0.0001$) and HCP-YA (rho = 0.29, $P_{perm} < 0.0001$) datasets. The developmental effects of the variability axis slope were still significant in the HCP-D dataset (partial $R^2 = -0.26$, $P = 4.23\times10^{-7}$, **Fig. S3g**). The cognitive effect of the variability axis showed a trend toward significance in the HCP-D dataset ($t = 1.85$, $P = 0.07$, partial $R^2 = 0.01$, **Fig. S3h**) and was significant in the HCP-YA dataset ($t = 3.15$, $P = 0.003$, partial $R^2 = 0.21$, **Fig. S3i**).

In addition, considering that the global signal exhibits network-specific effects and varies across development (47, 48), we assessed the robustness of our findings without applying global signal regression. As shown in **Fig. S2g,h**, the pattern of individual FC variability remained significantly correlated with structural communicability variability across connectome edges (HCP-D: rho = 0.31, $P_{perm} < 0.0001$; HCP-YA: rho = 0.27, $P_{perm} < 0.0001$). Consistently, we observed that the connectional axis slope of FC variability was significantly associated with age in youth (partial $R^2 = -0.59$, $P = 3.92\times10^{-26}$, **Fig. S3j**). The connectional axis slope of FC variability remained significantly associated with the fluid cognition composite score in both the HCP-D ($t = 2.60$, $P =$

0.012, partial $R^2$ = 0.06, **Fig. S3k**) and HCP-YA ($t$ = 12.23, $P = 4.28\times10^{-15}$, partial $R^2$ = 0.75, **Fig. S3l**) datasets.

As for the association between individual FC variability and structural communicability variability, we further applied a minimum tract threshold within participants to minimize the influence of spurious structural connections. A range of density levels (5% to 25%) was tested, and the structural communicability variability, estimated from binarized structural connectomes, remained significantly correlated with FC variability across these density levels in both the HCP-D and HCP-YA datasets (**Supplementary Table S1**).

Finally, we have also conducted sensitivity tests to validate our sliding window configurations in developmental and cognitive analyses by varying the window lengths (40, 50, and 60 participants) and step sizes (5 and 10 participants). Both the developmental (**Supplementary Table S2**) and cognitive (**Supplementary Table S3**) results remained relatively stable across the different combinations of window lengths and step sizes, confirming the robustness of our findings. The cognitive results of the HCP-D dataset were not significant when each window included only 40 participants (window length). However, the results became significant with increased window length, highlighting the importance of an adequate number of participants for reliable estimation of individual variability.

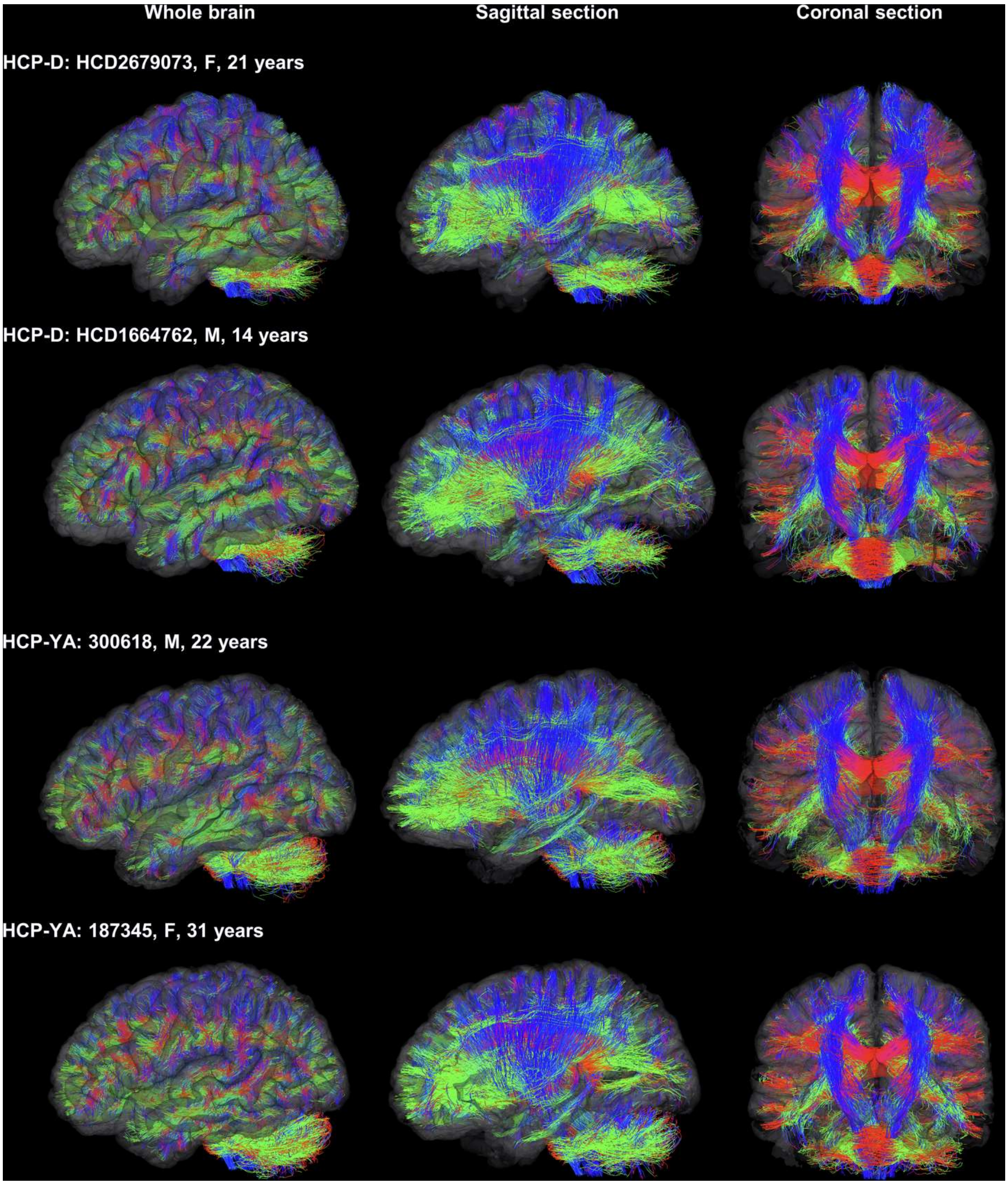


**Fig. S1. Exemplar visualization of white matter fiber tracts from probabilistic tractography**. Data from two randomly selected participants in the HCP-D dataset and two from the HCP-YA dataset are shown. The streamlines were generated using the probabilistic iFOD2 algorithm. Whole-brain view: the whole-brain fibers were loaded onto translucent cortical pial surfaces; Sagittal section: a clipping plane at the inferior frontal sulcus reveals key structures such as the corona radiata (blue bundles), anterior and posterior limbs of the internal capsule (green bundles), and a portion of the inferior longitudinal fasciculus within the temporal lobe; Coronal section: a clipping plane near the central sulcus highlights the corona radiata (blue bundles) and corpus callosum (red bundles). The colors represent fiber orientations: red for left-right, blue for superior-inferior, and green for anterior-posterior directions.

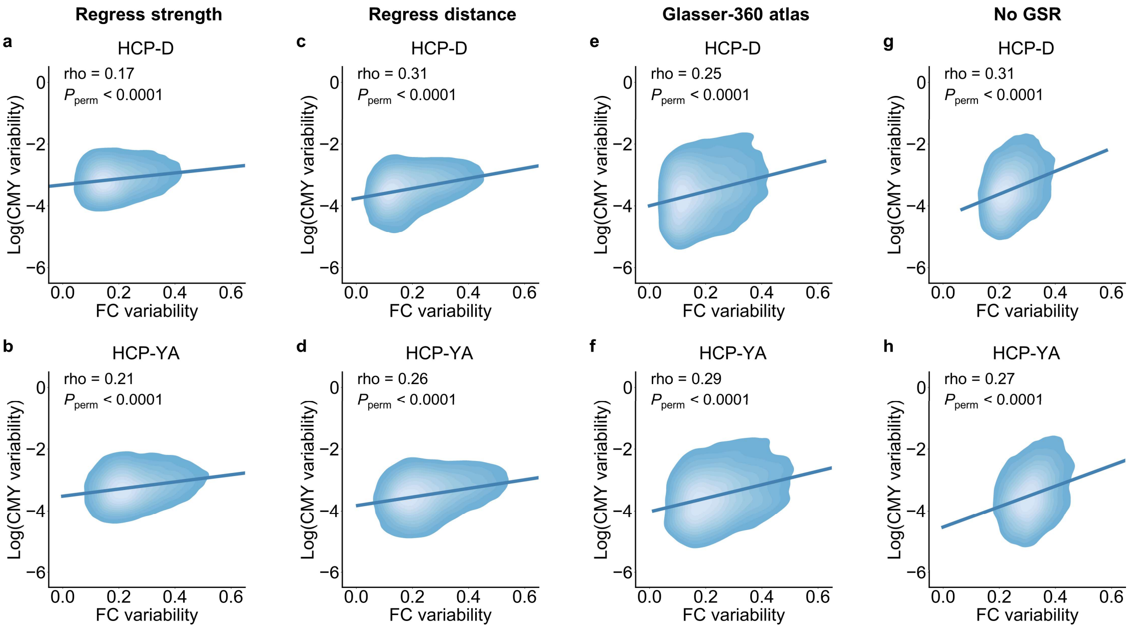


**Fig. S2. Sensitivity analyses of alignment between FC variability and SC variability.** FC variability was still positively correlated with structural communicability variability after 1) regressing out mean FC strength and structural communicability strength (**a**, rho = 0.17, HCP-D; **b**, rho = 0.21, HCP-YA); 2) regressing out inter-regional Euclidean distance (**c**, rho = 0.31, HCP-D; **d**, rho = 0.26, HCP-YA); 3) replicating with Glasser-360 atlas (**e**, rho = 0.25, HCP-D; **f**, rho = 0.29, HCP-YA); and 4) data pre-processing without global signal regression (**g**, rho = 0.31, HCP-D; **h**, rho = 0.27, HCP-YA). Spearman's rank correlations were used and all correlations were significant ($P_{perm}$ < 0.0001). CMY, communicability, GSR, global signal regression.

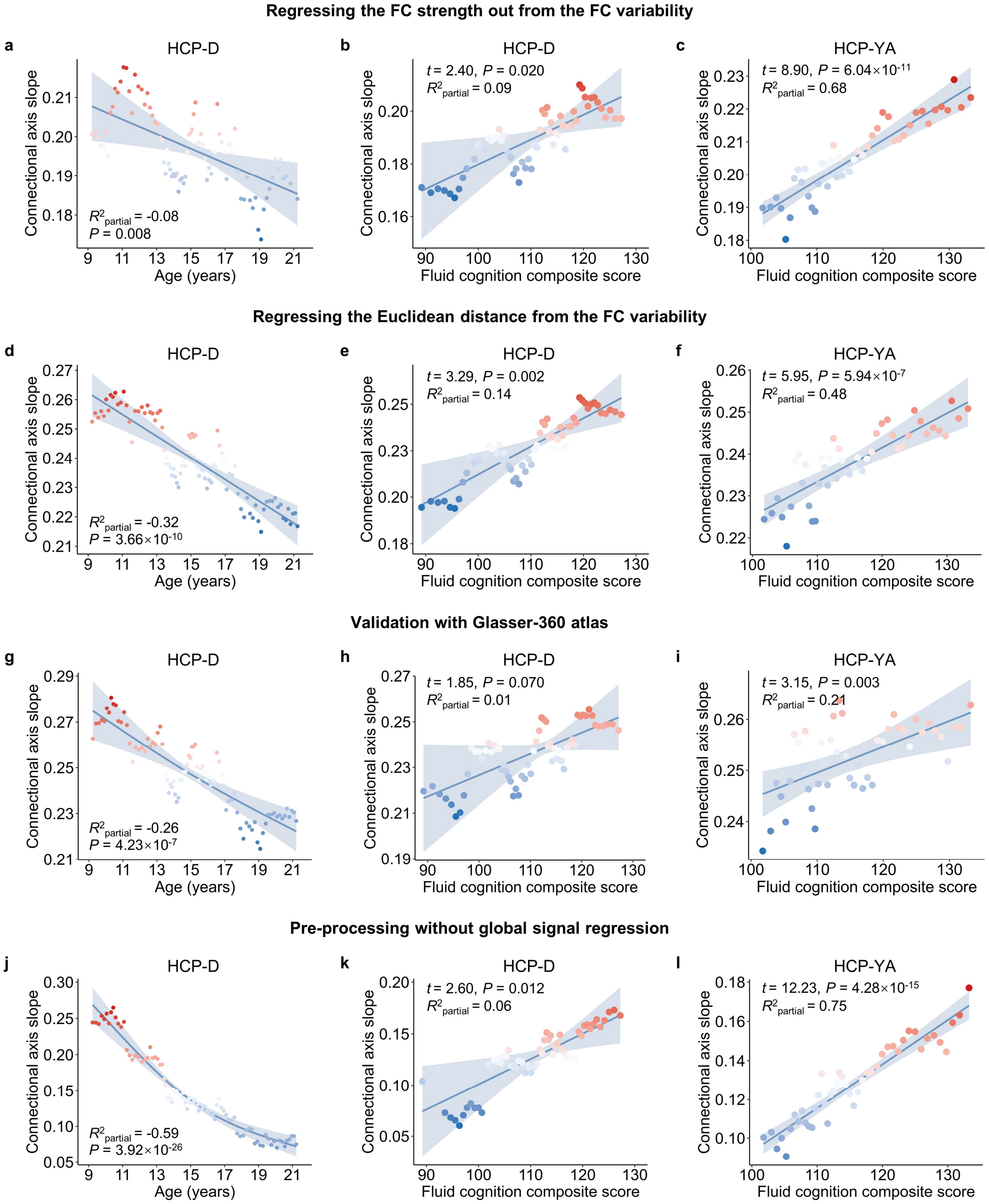


**Fig. S3. Sensitivity analyses of the developmental and cognitive effects of connectional variability axis.** After regressing out the FC strength from the individual FC variability of each group, the connectional axis slope of FC variability remained significantly declined with age in youth (**a**, partial $R^2$ = -0.08, $P$ = 0.008), and the connectional axis slope of FC variability remained significantly associated with the fluid cognition composite score in both the HCP-D (**b**, $t$ = 2.40, $P$ = 0.02, partial $R^2$ = 0.09) and HCP-YA (**c**, $t$ = 8.90, $P$ = 6.04×10$^{-11}$, partial $R^2$ = 0.68) datasets. Similarly, after regressing out the inter-regional Euclidean distance from the individual FC variability of each group, the connectional axis slope of FC variability remained significantly declined with age in youth (**d**, partial $R^2$ = -0.32, $P$ = 3.66×10$^{-10}$), and the connectional axis slope of FC variability remained significantly associated with the fluid cognition composite score in both the HCP-D (**e**, $t$ = 3.29, $P$ = 0.002, partial $R^2$ = 0.14) and HCP-YA (**f**, $t$ = 5.95, $P$ = 5.94×10$^{-7}$, partial $R^2$ = 0.48) datasets. Then, using the Glasser-360 atlas for connectivity calculation, the connectional axis slope of FC variability remained significantly declined with age in youth (**g**, partial $R^2$ = -0.26, $P$ = 4.23×10$^{-7}$), and the association between connectional axis slope of FC variability and the fluid cognition composite score showed a trend toward significance in the HCP-D (**h**, $t$ = 1.85, $P$ = 0.07, partial $R^2$ = 0.01) dataset and was significant in the HCP-YA (**i**, $t$ = 3.15, $P$ = 0.003, partial $R^2$ = 0.21) dataset. Finally, when re-processing the data without global signal regression, the connectional axis slope of FC variability remained significantly declined with age in youth (**j**, partial $R^2$ = -0.59, $P$ = 3.92×10$^{-26}$), and the connectional axis slope of FC variability remained significantly associated with the fluid cognition composite score in both the HCP-D (**k**, $t$ = 2.60, $P$ = 0.012, partial $R^2$ = 0.06) and HCP-YA (**l**, $t$ = 12.23, $P$ = 4.28×10$^{-15}$, partial $R^2$ = 0.75) datasets.

**Table S1. Association between FC variability and SC variability with binarized structural connectome.** The structural connectome (SC) was first binarized within each individual using the same threshold (density: 5% to 25%). Structural communicability variability was then calculated based on the binarized SC. Spearman's rank correlation was applied to estimate the association between FC variability and SC variability.

| | Spearman's rho | $P_{perm}$ |
|---|---|---|
| **HCP-D** | | |
| Density = 5% | 0.26 | < 0.0001 |
| Density = 10% | 0.28 | < 0.0001 |
| Density = 15% | 0.27 | < 0.0001 |
| Density = 20% | 0.25 | < 0.0001 |
| Density = 25% | 0.23 | < 0.0001 |
| **HCP-YA** | | |
| Density = 5% | 0.24 | < 0.0001 |
| Density = 10% | 0.25 | < 0.0001 |
| Density = 15% | 0.23 | < 0.0001 |
| Density = 20% | 0.22 | < 0.0001 |
| Density = 25% | 0.20 | < 0.0001 |

**Table S2. Developmental effects.** The developmental effects of connectional axis slope of FC variability with varying sliding window parameters (window length, step size) in the HCP-D dataset.

| Developmental effects | HCP-D | |
|---|---|---|
| | partial $R^2$ | $P$ |
| **Step = 5** | | |
| Length = 40 | -0.25 | $6.67\times10^{-8}$ |
| Length = 50 | -0.33 | $1.80\times10^{-10}$ |
| Length = 60 | -0.35 | $1.97\times10^{-11}$ |
| **Step = 10** | | |
| Length = 40 | -0.25 | 0.0002 |
| Length = 50 | -0.33 | $9.09\times10^{-6}$ |
| Length = 60 | -0.35 | $5.74\times10^{-6}$ |

**Table S3. Cognitive effects.** Association between connectional axis slope of FC variability and fluid cognition composite score with varying sliding window parameters (window length, step size) in the HCP-D and HCP-YA datasets.

| Cognitive effects | HCP-D | | | HCP-YA | | |
|---|---|---|---|---|---|---|
| | *t* | *P* | partial $R^2$ | *t* | *P* | partial $R^2$ |
| **Step = 5** | | | | | | |
| Length = 40 | 1.63 | 0.1086 | 0.04 | 5.05 | $9.25\times10^{-6}$ | 0.39 |
| Length = 50 | 3.53 | 0.0008 | 0.15 | 5.08 | $9.62\times10^{-6}$ | 0.41 |
| Length = 60 | 5.01 | $6.08\times10^{-6}$ | 0.25 | 6.11 | $4.23\times10^{-7}$ | 0.50 |
| **Step = 10** | | | | | | |
| Length = 40 | 0.81 | 0.4262 | 0.02 | 2.97 | 0.0079 | 0.32 |
| Length = 50 | 2.09 | 0.0465 | 0.14 | 3.09 | 0.0063 | 0.35 |
| Length = 60 | 2.97 | 0.0065 | 0.19 | 4.80 | 0.0002 | 0.51 |
